# Infectious bronchitis virus attaches to lipid rafts and enters cells via clathrin mediated endocytosis

**DOI:** 10.1101/352898

**Authors:** Huan Wang, Yingjie Sun, Xiang Mao, Chunchun Meng, Lei Tan, Cuiping Song, Xusheng Qiu, Chan Ding, Ying Liao

**Author notes:** Corresponding author: Ying Liao, Chan Ding. These authors contributed equally as first author (Huan Wang, Ying Liao).

## Abstract

Due to its economic importance to in poultry industry, the biology and pathogenesis of infectious bronchitis virus (IBV) have been investigated extensively. However, the molecular mechanisms involved in IBV entry are not well characterized. In this study, systematic approaches were used to dissect IBV entry process in various susceptible cells. First, we observed that lipid rafts were involved in IBV attachment. Second, low pH in intracyplasmic vesicles was required for virus entry. By using the specific clathrin mediated endocytosis (CME) inhibitor or knock down of clathrin heavy chain (CHC), we demonstrated that IBV mainly utilized the CME for its entry. Furthermore, GTPase dynamin1 was involved in virus containing vesicle scission and internalization. Surprisingly, CME adaptor Eps15 had no effect on IBV internalization. Third, the penetration of IBV into cells led to active cytoskeleton rearrangement. After internalization, virus particles moved along with the classical endosome/lysosome track, as evidenced by co-localization of R18 labeled IBV with vehicle markers Rab5/Rab7/LAMP1 along with the infection time course. Functional inactivation of Rab5 and Rab7 significantly inhibited IBV infection. VCP, a protein helps early endosome maturation, was involved virus trafficking. Finally, by using the dual R18/DiOC labeled IBV, we observed that membrane fusion with late endosome/lysosome membranes was induced between 2-3 h.p.i.. Taken together, our findings demonstrate that IBV virions attach to lipid rafts and are internalized into cells via CME, move along with early/late endosomes-lysosomes, finally fuse with late endosome-lysosome membranes, release virus genome into cytoplasm. This study provides comprehensive images of IBV attachment-internalization-trafficking-fusion steps.

**IMPORTANCE:** IBV, the avian coronavirus isolated in 1937, infects chicken and causes economic loss in poultry industry. It has been reported that the entry of IBV requires low pH. However, the molecular mechanisms underlying IBV internalization and trafficking remain to be clarified. Therefore, we employed multiple chemical and molecular approaches to dissect the entry mechanisms of IBV in susceptible cells. Our results showed IBV entry was significantly inhibited when clathrin-mediated endocytosis (CME) was blocked by chemical inhibitor or depletion of clathrin protein. Moreover, by using R18-labeled IBV, we found that IBV particles attached to lipid rafts, led to actin rearrangement, and moved along with the entire endosomal system. R18/DiOC labeling method showed that IBV fused with late endosomes or lysosomes. This is the first report to describe the entire entry process of IBV, allowing for a better understanding of the infection process of group III avian coronavirus.

## INTRODUCTION

Most viruses take advantage of cellular endocytic pathways to enter their host cells (1, 2). Once internalized, virus particles move through a dynamic network of endocytic vesicles, which undergoes gradual sorting and complex maturation events. Endosome maturation, in turn, triggers conformational changes and dissociation events in the incoming viruses, which ultimately lead to the delivery of the viral genome and associated proteins into the cytoplasm. In general, while non-enveloped viruses are able to penetrate the limiting endosomal membrane by lysis or pore formation (3), enveloped viruses fuse with the vesicle membranes to release the genome into the cytoplasm (4). Different viruses employ different routes of endocytosis, and the route taken by a given virus largely depends on the receptor that the virus interacts with. Numerous virus families utilize endocytosis to infect host cells, mediating virus internalization as well as trafficking to the site of replication. The endocytic pathways utilized by viruses include clathrin-mediated endocytosis (CME), caveolin/raft-mediated endocytosis (CavME), macropinocytosis, and non-clathrin/non-caveolin mediated endocytosis. CME is the best understood endocytic pathway. In response to receptor mediated internalization signals, clathrin is recruited to plasma membrane, leading to the assembly of clathrin-coated pits (CCP) at the cytoplasmic side of the plasma membrane. The main scaffold component of clathrin coat is the clathrin heavy chain (CHC) and clathrin light chain (CLC), forming a three-legged trimers called triskelions (5). The adaptor protein 2 (AP-2) provides a bridge between receptor cargo domain and the clathrin β subunit (6). Once assembled, CCPs pinch off from the plasma membrane and mature into clathrin-coated vesicles (5). During this process, the adaptors also play an important role, including Eps15 and dynamin GTPase. Lots of viruses use CME to enter cells, including Semliki forest virus, vesicular stomatitis virus (VSV) (7), hepatitis C virus, and Adenoviruses 2 and 5 (8). The Caveolae is a relatively small vesicle with a diameter of 50 to 100 nm, which is a flask-shaped invasion formed by membrane retraction on the plasma membrane and is a specialized lipid raft domain composed of caveolin and high levels of cholesterol (9, 10). CavME is first observed for the cellular uptake of simian virus 40 (11). After that, other viruses including some of the picornaviruses (12), papillomaviruses (13), filoviruses (14), murine leukemia virus (15), and human coronavirus 229E (HCoV-229E) (16) are found by utilizing CavME to enter into cells. Macropinocytosis is morphologically defined by the presence of membranous extensions of outwardly polymerizing actin termed membrane ruffles. It has been identified as an entry mechanism for several pathogens, including B2 adenovirus (17), Coxsackie B virus (18), influenza A virus (19), Ebola virus (20), Vaccinia virus (21), Nipah virus (22), Kaposi’s Sarcoma-Associated Herpesvirus (23), Newcastle disease virus (24).

Lipid rafts are functional membrane microdomains which are also termed detergent-insoluble glycolipid-enriched complexes or detergent-resistant membranes, rich in cholesterol, sphingolipids, and proteins (25, 26). They serve as domains to concentrate membrane-associated proteins that include receptors and signaling molecules (25). In addition to physiological roles in signal transduction of host cells (27), lipid rafts often serve as a site for entry, assembly and budding of microbial pathogens (28-30). They are involved in the binding and entry of host cells for several enveloped and non-enveloped viruses, including human immunodeficiency virus type 1 (31), poliovirus (32), human herpes virus 6 (33), West Nile virus (34), foot-and-mouth disease virus (35), and dengue virus(36). There is some evidence that lipid rafts are also involved in coronavirus entry. In the case of HCoV-229E, virus entry is inhibited by depletion of cholesterol, which results in the disruption of viral association with the cellular receptor CD13 (16). Thorp and Gallagher show that lipid rafts are crucial for the entry of mouse hepatitis virus (MHV) (37). Guo et al., demonstrate that lipid rafts are involved in IBV entry (38).

Coronaviruses are enveloped, plus-strand RNA viruses belonging to the family *Coronaviridae*, which includes many human and animal pathogens of global concern, such as severe acute respiratory syndrome coronavirus (SARS-CoV), middle east respiratory syndrome virus (MERS-CoV), HCoV-229E, MHV, porcine epidemic diarrhea virus (PEDV), Feline infectious peritonitis virus (FIPV), and IBV. In most cases, they cause respiratory and/or intestinal tract disease. In general, coronaviruses enter cells by endocytosis, which undergoes pH-dependent membrane fusion. For example, CME as well as clathrin- and caveolae-independent entry pathways have been reported for SARS-CoV (39, 40); clathrin- and serine proteases-dependent uptake has been reported for PEDV (41); FIPV is suggested to enter cells via a clathrin- and caveolae-independent endocytic route (42); and MHV2 enters cells via CME, but independent of Eps15 (43). IBV belongs to the genus *gamma coronavirus*. Its genome is approximately 27.6 kilobases (kb) in length, encoding 2 polyproteins (1a, 1ab), four structural proteins, namely, spike protein (S), membrane protein (M), and small envelope protein (E), nucleocapsid protein (N); and four accessory proteins named 3a, 3b, 5a, 5b, are encoded by subgenomic RNA species. Polyprotein 1a and 1ab are cleaved by viral proteases into at least 15 non-structural proteins (nsp2-nsp16). The structural and accessory proteins are encoded by subgenomic RNA species. Despite the importance of IBV as one of the dominant pathogens causing a highly contagious infectious bronchitis worldwide and leading to significant economic loss in the chicken industry, little information is available about its entry mechanisms. It has been identified that the entry of IBV into cells depends on lipid rafts and low pH (44); however, the endocytic pathway that IBV hijacks is unclear.

The early step in the entry process of IBV into target cells is initiated by engagement of the S glycoprotein with the receptor. Here, using several cell lines permissive to IBV Beaudette strain, we report that IBV virion attaches to lipid rafts in the cell surface and the efficient entry of IBV into cells requires the presence of intact lipid rafts. We also address IBV entry pathway by systematically perturbing the function of key factors in the various endocytic mechanisms using chemical inhibitors, siRNA silence, and overexpression of dominant negative proteins. To achieve a successful tracking of virion attachment, entry, intracellular trafficking, and membrane fusion, octadecyl rhodamine (R18) and 3, 3′-Dihexyloxacarbocyanine Iodide (DiOC) were labeled on viral membrane as fluorescent tags. Immunofluorescence analysis was used to visualize the co-localization or co-trafficking of virus and cellular factors. Our results reveal that IBV is internalized into cells by CME, transports along early endosomes, late endosomes, and fuses with membrane at late endosome-lysosomes.

## MATERIALS AND METHODS

### Cells and viruses

Vero cells, Huh7 cells, BHK-21, and DF-1 cells were maintained in Dulbecco’s modified Eagle medium (DMEM) with 4500 mg/l glucose, supplemented with 10% fetal bovine serum (FBS) (Hyclone, USA), 100 units/ml penicillin and 100 μg/ml streptomycin (Invitrogen, USA). H1299 cells were maintained in RPMI 1640, supplemented with 10% FBS in the presence of penicillin and streptomycin. Above cells were purchased from ATCC (USA) and cultured at 37°C with 5% CO_2_.

The Beaudette strain of IBV (ATCC VR-22) adapted to Vero cells was used in this study. As a well characterized virus, Vesicular Stomatitis Virus (VSV) was used as control virus. Virus stock was prepared by infecting monolayers of Vero cells with multiplicity of infection (MOI) of 0.1 in FBS free DMEM. After attachment for 1 hour (h), the unbounded virus was removed and replaced with FBS free DMEM. The virus and cells were incubated at 37°C until 100% cells were with cytopathic effect (CPE). After freezing and thawing the virus containing liquid for three times, cell debris was removed by centrifugation at 3000×rpm, and supernatant was aliquoted and stored at −80°C as virus stock. A control of Vero cell lysates from mock-infected cells was prepared in the same manner. The virus yield was assessed by titration on Vero cells in 96-well plates according to the method described by Reed and Muench(45).

### R18 and R18/DiOC labeling of IBV or VSV

Vero cells were infected with 0.1 MOI of virus until 100% cells were with CPE. The cell supernatant was clarified by centrifuged at 4000×g for 15 min. The supernatant was then centrifuged at 5000×g for 60 min and concentrated by 100-fold by using Amicon^®^ Ultra-15 Centrifugal Filter Devices (10-kDa cutoff, Merk, Poland) which provided fast ultrafiltration. Mock infected Vero cells culture medium was concentrated in the same manner as control sample in labeling experiment.

R18 labeling was prepared as described previously (44): 100 μl of concentrated virus or control sample was incubated with 2.5 μl of 1.7 mM R18 (Molecular Probes, USA) on a rotary shaker for 1 h at room temperature (in the dark). R18/DiOC labeling was prepared as described (46): 100 μl of concentrated virus or control sample was incubated with 3.3 mM DiOC (Molecular Probes, USA) and 6.7 mM R18 mixture (Molecular Probes, USA) with gentle shaking for 1 h at room temperature. After finishing labeling, the virus and dye mixture was re-suspended in 8 ml phosphate-buffered saline (PBS), and the excess dye was removed with an Amicon^®^ Ultra-15 Centrifugal Filter Devices (10-kDa cutoff, Merk, Poland) by centrifugation for 60 min. Finally, 100 μl of R18 or R18/DiOC labeled-virus were obtained.

### Inhibitors and antibodies

The endocytotic pathway inhibitors Amiloride (S1811), Nystatin (S1934), and chlorpromazine (CPZ, S2456) were purchased from Selleck (USA). Actin monomer-sequestering drug cytochalasin D (CytoD, PHZ1063), actin polymer-stabilizing jasplakinolide (Jas, J7473) were purchased from Thermo Fisher Scientific (USA). Endosome acidification inhibitor NH_4_Cl (A9434), cholesterol depletion drug methyl-β-cyclodextrin (MβCD, C4555) and Cholesterol-Water Soluble (C4951) were purchased from Sigma-Aldrich (USA).

Anti-IBV S and N antibodies were obtained through immunization of rabbits with respective antigens. Anti-flotillin-1 (#18634), anti-GFP (#2956), anti-CHC (clathrin heavy chain) (#4769s), anti-Rab5 (#3547s), anti-Rab7 (#9367s), and anti-LAMP1 (#9091s) were purchased from Cell Signaling Technology (USA). Anti-VSV G (Ab50549) was obtained from Abcam (UK). Anti-β-actin (A1978) was purchased from Sigma Aldrich (USA). Cholera Toxin Subunit B (CTB, C34775) was purchased from Thermo Fisher Scientific (USA). Fluorescein isothiocyanate (FITC)-conjugated anti-mouse or anti-rabbit immunoglobulin G (IgG), as well as horseradish peroxidase (HRP)-conjugated anti-mouse or anti-rabbit IgG were purchased from Cell Signaling Technology (USA). Goat anti-Rabbit IgG (H+L) Highly Cross-Adsorbed Secondary Antibody, Alexa Fluor 633 (A-21071) was purchased from Thermo fisher (USA)

### Virus infection and drug treatment

To test the effect of various inhibitors on IBV infection, Vero cells, H1299 cells, BHK-21 cells, or Huh7 cells were seeded into 6-well plates at 5×10^6^ cells/well and cultured for 24 h until they reached 100% confluence. Cells were then pretreated with the indicated concentrations of NH_4_Cl, CPZ, Nystatin, or Amiloride for 30 min at 37°C, respectively. After treatment, 1 MOI of virus was inoculated into each well and incubated at 4°C for 1 h in the absence of compounds. The unbound virions were washed away with cold PBS and the cells incubated at 37°C with respective compounds in fresh medium for 1 h. After that, cells were replenished with fresh medium without compounds. Virus internalization was determined by semi-quantitative real time RT-PCR at 2 h.p.i. by measuring the viral RNA genome, and virus replication was monitored by Western blot at 8 h.p.i. by checking the expression level of viral N protein.

### Cell viability assay and pH assessment

Viability of drug-treated cells was measured using the WST-1 Cell proliferation and cytotoxicity assay kit (Beyotime, Haimen, China) according to the manufacturer’s instructions. Cells were seeded in 96-well plate and treated with indicated drugs, 10 μl of WST-1 was added to each well and incubated for 1 h. The absorbance at 450 nm was monitored and the reference wavelength was set at 630 nm. The viability of cells was calculated by comparison to that of untreated cells.

To assess the effect of NH_4_Cl on the pH change of acidic intracellular vesicles, Vero cells were treated with increasing concentrations of NH_4_Cl for 30 min at 37°C, followed by staining with 1 mg/ml acridine orange in serum free medium for 15 min at 37°C. The cells were washed twice with PBS and stained with DAPI (2-(4-Amidinophenyl)-6-indolecarbamidine dihydrochloride, Roche). Images were taken with a Zeiss Axio Observer Z1 fluorescence microscope.

### Cholesterol depletion and replenishment

Cells were incubated with 5 mM MβCD for 30 min at 37°C at indicated infection time course. The cells exposed to DMSO for 30 min were set as control group. After incubation, the cells were washed twice with PBS and replaced with fresh culture medium. For cholesterol replenishment, cells were pretreated with 2.5 or 5 mM MβCD for 30 min at 37°C, then supplemented with or without exogenous 1 mM cholesterol and incubated for 1 h at 37°C. After extensive washing with PBS, the cells were subjected to IBV infection.

### Membrane flotation analysis

Cells (5 × 10^7^) were incubated with IBV (MOI of 5) at 4°C for 1 h, or mock-infected. Cells were lysed on ice for 30 min in 1 ml TNE lysis buffer with 1% Triton X-100, supplemented with complete protease inhibitor cocktail. The cell lysates were centrifuged at 4000×g for 5 min at 4°C, to remove cell debris and nuclei. The supernatant was collected and mixed with 1 ml of TNE buffer with 80% sucrose, placed at the bottom of the ultracentrifuge tube, and overlaid with 6 ml of 30% and 3 ml of 5% sucrose in TNE buffer. The lysates were ultracentrifuged at 38000×rpm for 18 h at 4°C in an SW41 rotor (Beckman). After centrifugation, 12 fractions (1 ml each) were collected from the top to the bottom of the tube and subjected to 8% or 10% SDS-PAGE, followed by Western blotting using antibodies of anti-Flotillin-1, anti-N protein, or anti-S protein.

### Plasmid transfection and siRNA transfection

Cells grown in 6-well plates were transfected with 3 μg of plasmid DNA, by using lipofectamine 3000 (Invitrogen) according to the manufacturer’s instructions. After 24 h post-transfection, cells were infected with IBV at MOI of 1. The internalization level of IBV genomic RNA was determined by semi-quantitative real time RT-PCR at 2 h.p.i., the levels of viral proteins expression were measured by Western blot analysis at 8 h.p.i., and the virus particle release in the supernatant was assessed by TCID_50_ measurement at 12 h.p.i..

To knock down Eps15, dynamin1, Rab5, or Rab7, cells were seeded into 6-well plates, and small interfering RNA (siRNA) duplexes at a concentration of 100 nM were transfected into the cells by using Lipofectamine 3000 according to the manufacturer’s instructions. At 36 h post-transfection, cells were infected with IBV at MOI of 1. The level of IBV internalization, protein expression, and virus particles release were then measured by semi-quantitative real time RT-PCR, Western blot analysis, and TCID_50_ at indicated time points. The siRNA sequences were shown in table 1.

**Table1.**
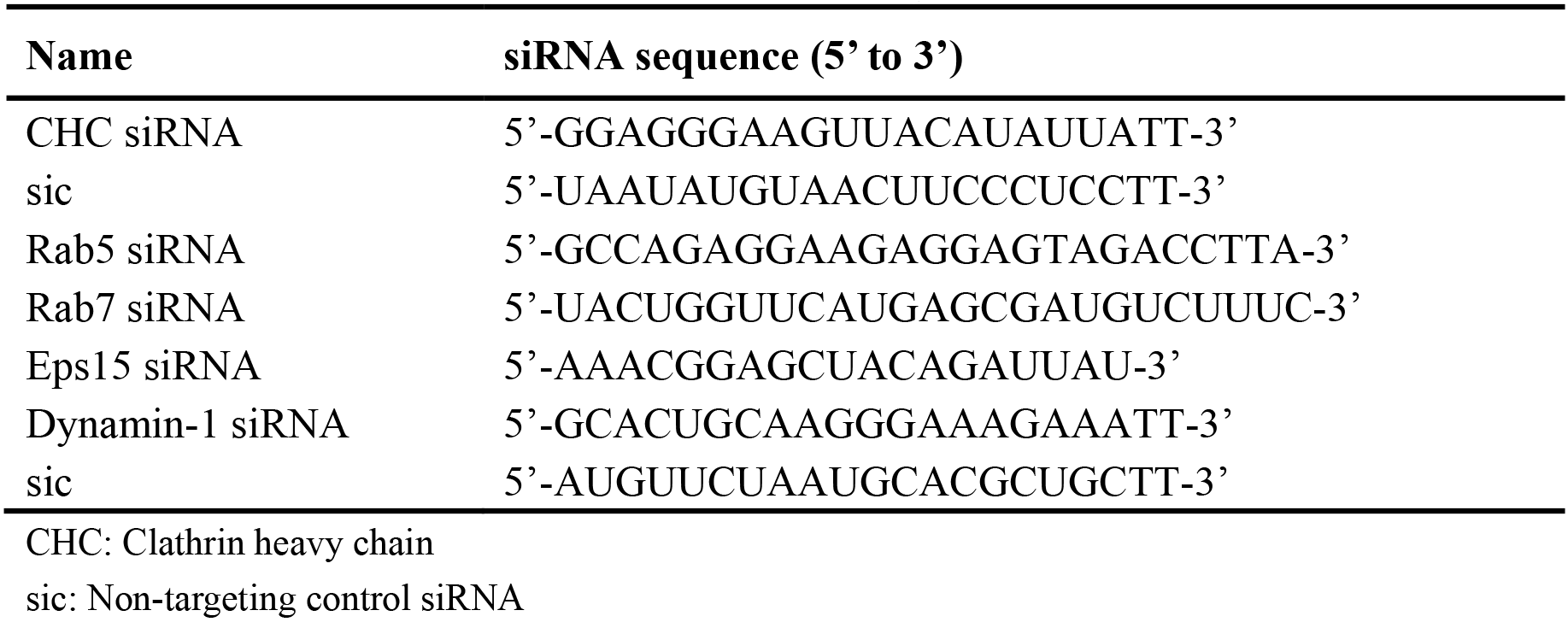
siRNA Sequence.

### Measurement of virus genomic RNA (gRNA) by semi-quantitative real time RT-PCR

Viral internalization was detected as described previously (47). Briefly, the cells in 6-well plates were incubated with IBV (MOI=1) at 4°C for 1 h. The unbound virions were washed away with cold PBS, and the cells were incubated at 37°C with fresh medium. At 2 h.p.i., cells were treated with 1 mg/ml proteinase K (Invitrogen) for 15 min to remove adsorbed but not internalized virus. Proteinase K was then inactivated with 2 mM phenylmethylsulfonyl fluoride (PMSF) in PBS with 3% bovine serum albumin. Cells were then washed three times with PBS and lysed in 1 ml of TRIZOL reagent (Invitrogen, USA) per well for RNA isolation. One fifth volume of chloroform was added. The mixture was centrifuged at 13,000×rpm for 15 10min at 4C, and the aqueous phase was then mixed with an equal volume of 100% isopropanol and incubated at room temperature for 10 min. RNA was precipitated by centrifugation at 13,000xrpm for 10 min at 4°C. RNA pellets were washed with 70% RNase-free ethanol and dissolved in RNase-free water.

The level of internalized viral genomic RNA (+gRNA) or negative genomic RNA (-gRNA) was determined by real time RT-PCR using primers targeting to IBV genome. Briefly, 3 μg of total RNA was used to perform reverse transcription using expand reverse transcriptase (Roche, USA) and oligo-dT/specific primers. Equal volume of cDNAs was then PCR-amplified using SYBR green PCR master kit (Dongsheng Biotech, Guangdong, China). Genome copy numbers were normalized to β-actin levels by using the comparative cycle threshold values determined in parallel experiment. Data were analyzed relative to controls. All assays were performed in three replicates. The primers sequence used in RT-PCR were shown in Table 2.

**Table 2.**
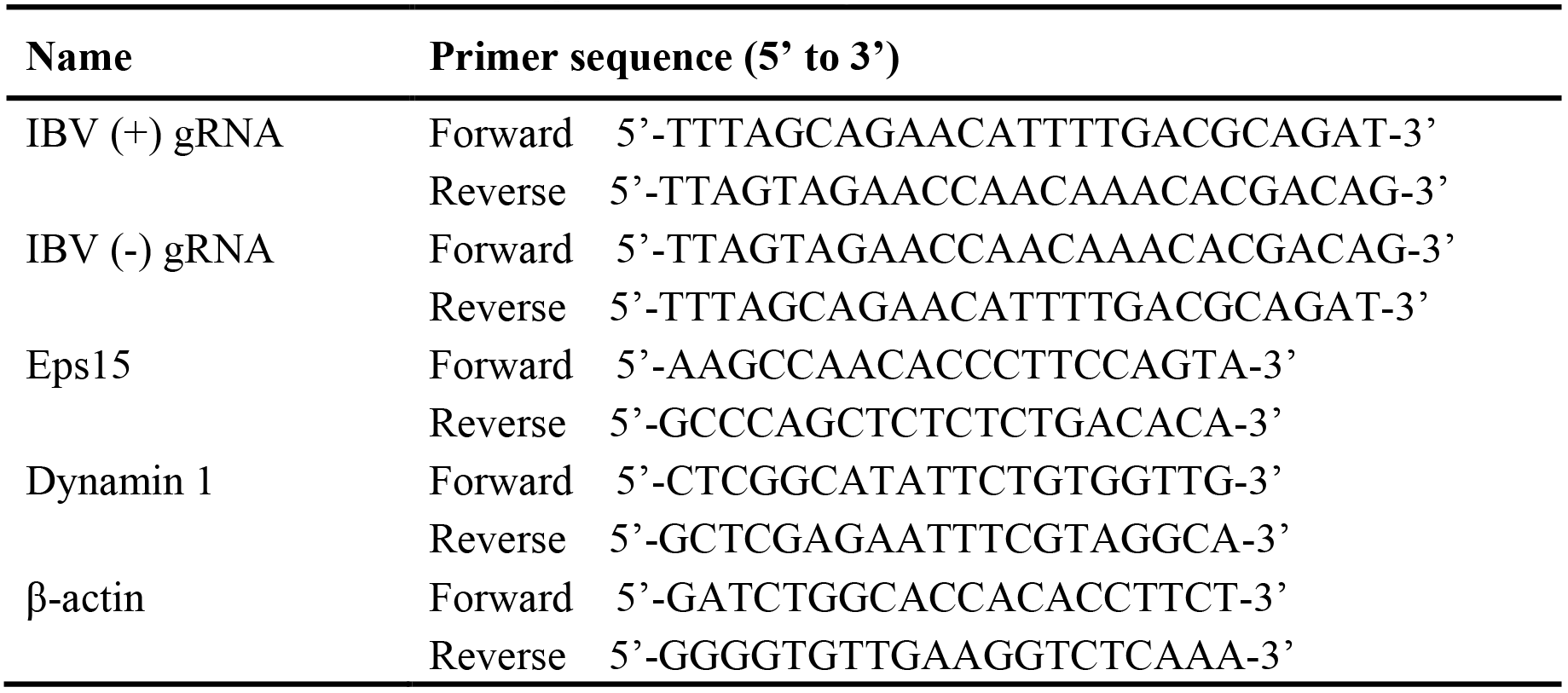
Primar sequence for semi-quantitative real time RT-PCR.

### Western blot analysis

Cells were lysed with 1x SDS loading buffer in the presence of 100 mM dithiothreitol and denatured at 100°C for 5 min. Equivalent amounts of protein were separated by SDS-PAGE, followed by transferring onto polyvinylidenedifluoride (PVDF) membranes (Bio-Rad Laboratories, USA) by electroblotting. Immunoblot analysis was then performed by incubating membranes with blocking buffer for 1 h at room temperature, and with appropriate antibodies diluted in blocking buffer for another 1 h. After washing three times with PBST, membranes were incubated with HRP-conjugated secondary antibody for 1 h and washed with PBST thrice. Blots were developed with an enhanced chemiluminescence (ECL) detection system (GE Healthcare Life Sciences, USA) and exposed to Chemiluminescence gel imaging system (Tanon 5200, Shanghai, China). Membranes were stripped with stripping buffer (10 mM β-mercaptoethanol, 2% SDS, 62.5 mM Tris-Cl, pH 6.8) at 55°C for 30 min before re-probing with other antibodies.

### Immunostaining

Cells were seeded on 4-well chamber slides and infected with R18-IBV or R18-DiOC-IBV at MOI of 5. At indicated time points, cells were fixed with 4% paraformaldehyde for 10 min, washed thrice with PBS, permeabilized with 0.2% Triton X-100 for 10 min, and washed thrice with PBS. For the detection of GM1 by CTB, no cell permeabilization required. Cells were then blocked in 3% FBS for 1 h, and incubated with anti-Rab5, anti-Rab7, anti-LAMP1, anti-phalloidin (1:200 diluted in PBS, 5% BSA), or CTB (5 μg/ml), for 2 h, washed thrice with PBS, and then incubated with secondary antibody conjugating with FITC (DAKO) for 2 h (1:200 diluted in PBS, 5% BSA), followed by PBS washing. Cells were incubated with 0.1 μg/ml DAPI for 10 min and rinsed with PBS. Finally, the specimen was mounted with glass cover slips using fluorescent mounting medium (DAKO) containing 15 mMNaN3. Images were collected with a LSM880 confocal laser-scanning microscope (Zeiss).

### Virus titration

Supernatants of IBV-infected cells collected at 12 h.p.i. were centrifuged at 10000×g for 15 min to remove cell debris. The viral titers were determined by 50% infectious dose (TCID_50_) assay. Briefly, 10-fold serially diluted aliquots of virus were applied to confluent monolayers of DF-1 cells in 96-well plates. After 1 h of absorption, unbound viruses were removed, and the cells were washed twice with DMEM. The plates were incubated with DMEM at 37°C and the cytopathic effect (CPE) was observed after 3 days. Each sample was titrated in triplicate. The tissue culture TCID_50_ is calculated using Reed and Munch mathematical analysis of the data (45).

### Plasmid construction

Eps15, Rab5 and Rab7 were PCR amplified from HeLa cellular cDNAs and cloned into vector pEGFPN1 between restrict enzyme *Xho I* and *BamH I* under control of a cytomegalovirus promoter, generating pEGFPN1-Eps15, pEGFPN1-Rab5, and pEGFPN1-Rab7 with GFP-tag at N-terminus. Rab5-DN (S34N), Rab5-CA (Q79L), and Rab7-CA (Q67L) mutants were constructed by site-directed overlapped two round PCR mutagenesis. Plasmid Rab7-DN (T22N) was constructed as described previously (21) using the TaKaRa Mutant BEST Kit (Takara Bio, Dalian, China, #R401). Valosin-Containing Protein (VCP) was PCR amplified from HeLa cellular cDNA and cloned into vector pEGFPN1 between restrict enzyme *Hind* ***III*** and *Kpn* ***I*** under control of a cytomegalovirus promoter, generating pEGFPN1-VCP with GFP-tag at N-terminus. The pEGFPN1-Eps15-DN, pEGFPN1-dynamin1-WT and pEGFPN1-dynamin1-DN (K44A) constructs were generous gifts from Prof. Mao Xiang. All primers and restriction enzymes used for plasmids construction are shown in Table 3.

**Table 3.**
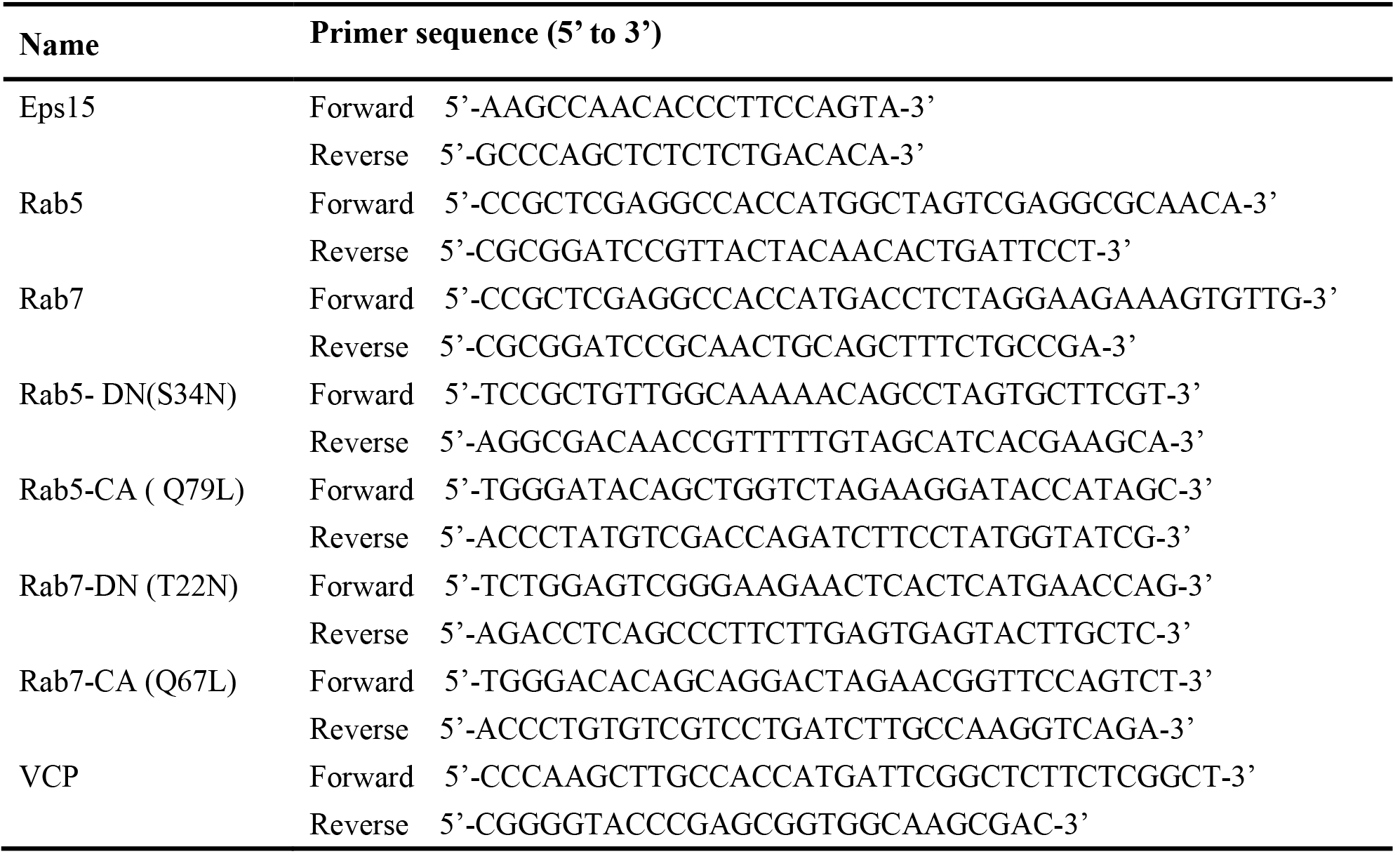
Primer sequence for plasmid construction.

### Statistical analysis

All data are presented as means ±standard deviations (SD), as indicated. Student’s t test was used to compare data from pairs of treated or untreated groups. Statistical significance is indicated in the figure legends. All statistical analyses and calculations were performed by using Graph Pad Prism 5 (Graph Pad Software Inc., La Jolla, CA).

### Densitometry

The intensities of corresponding bands were quantified using ImageJ program (NIH).

## RESULTS

### IBV entry is lipid raft-associated

Lipid rafts are cholesterol-enriched microdomains, where many cellular proteins, including viral receptors, are preferentially localized at. Various viruses enter host cells through lipid rafts (26). Previous report showed that IBV structural proteins migrate and integrate into lipid rafts during infection, while non-structural proteins do not integrate into rafts (38), suggesting that lipid rafts might participate in virus entry or budding/release. To examine which stage of infection that lipid rafts are involved in, MβCD was used to disrupt lipid rafts. The subtoxic dose of the MβCD in IBV permissive cell lines was determined by a cell proliferation assay (Supplementary Fig. 1). After determination of subtoxic dose, Vero, H1299, Huh7 cells were treated with 5 mM MβCD for 30 min at pre-, during-, or post-IBV infection at −0.5, 0, 1, 2 h.p.i. (Fig. 1A). The effect of MβCD on virus infection was accessed by checking viral gRNA, viral protein, and progeny virus with semi-quantitative real time RT-PCR, Western blot, and TCID_50_ assay, respectively. As shown in Fig. 1B, 1E, and 1H, treatment of various cells with MβCD at −1.5 and 0 h.p.i. acquired maximum blockage of virus genome level, which was determined by checking negative stranded genomic RNA at 4 h.p.i.. Surprisingly, treatment with MβCD at 1 and 2 h.p.i. did not affect the virus genome production. Next, viral N protein expression was analyzed with Western blot at 8 h.p.i.. As shown in Fig. 1C, 1F, 1I, treatment with MβCD −1.5 and 0 h.p.i. obtained maximum inhibition effect on viral N protein expression in various cell lines, whereas treatment with MβCD at post-entry step did not affect the viral N protein expression. In consistence, the progeny virus release was also inhibited by MβCD treatment at −1.5 and 0 h.p.i., but not by post-entry treatment (Fig. 1D, 1G; 1J). In all, above data demonstrate that MβCD inhibits virus entry step, but not post-entry process, suggesting that intact lipid rafts are required for virus attachment and/or entry.

**Figure 1.**
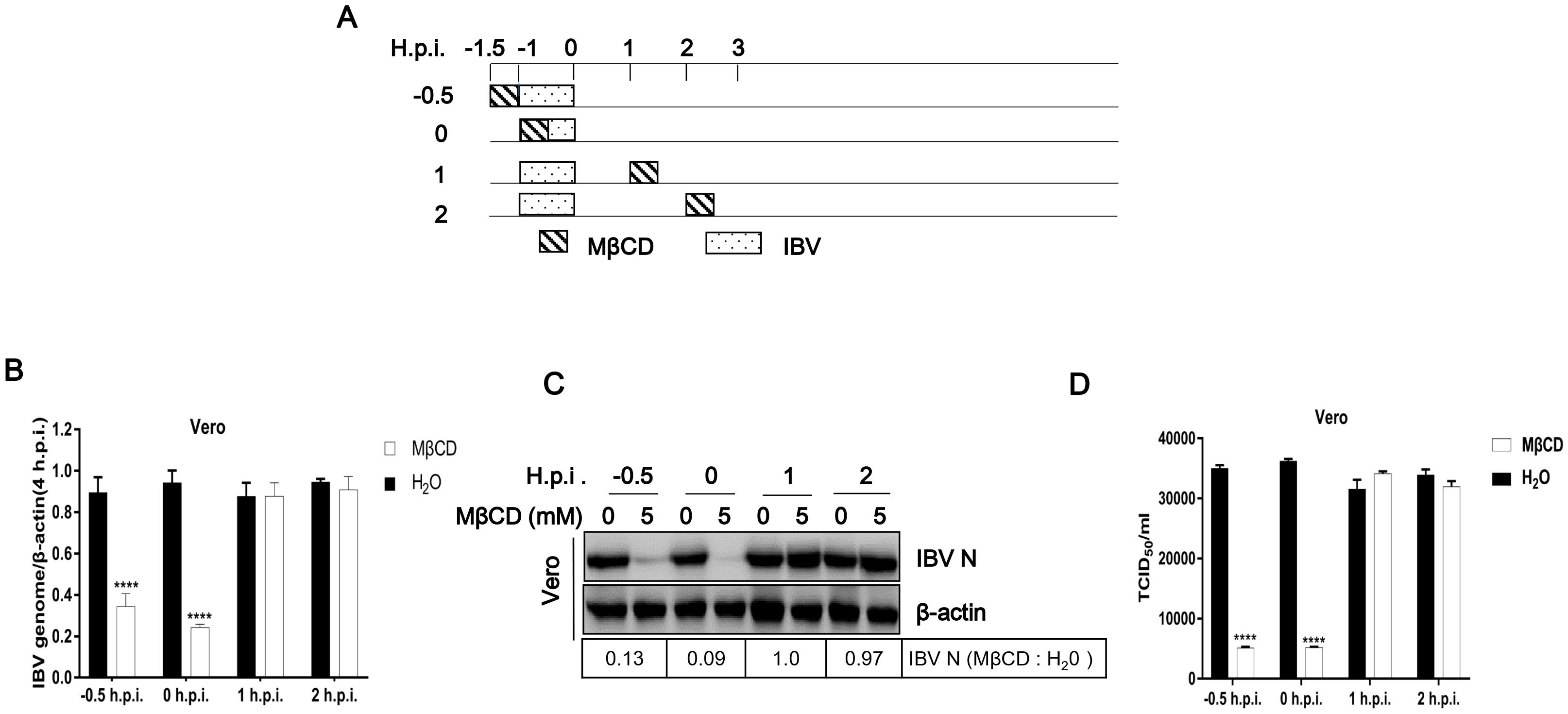
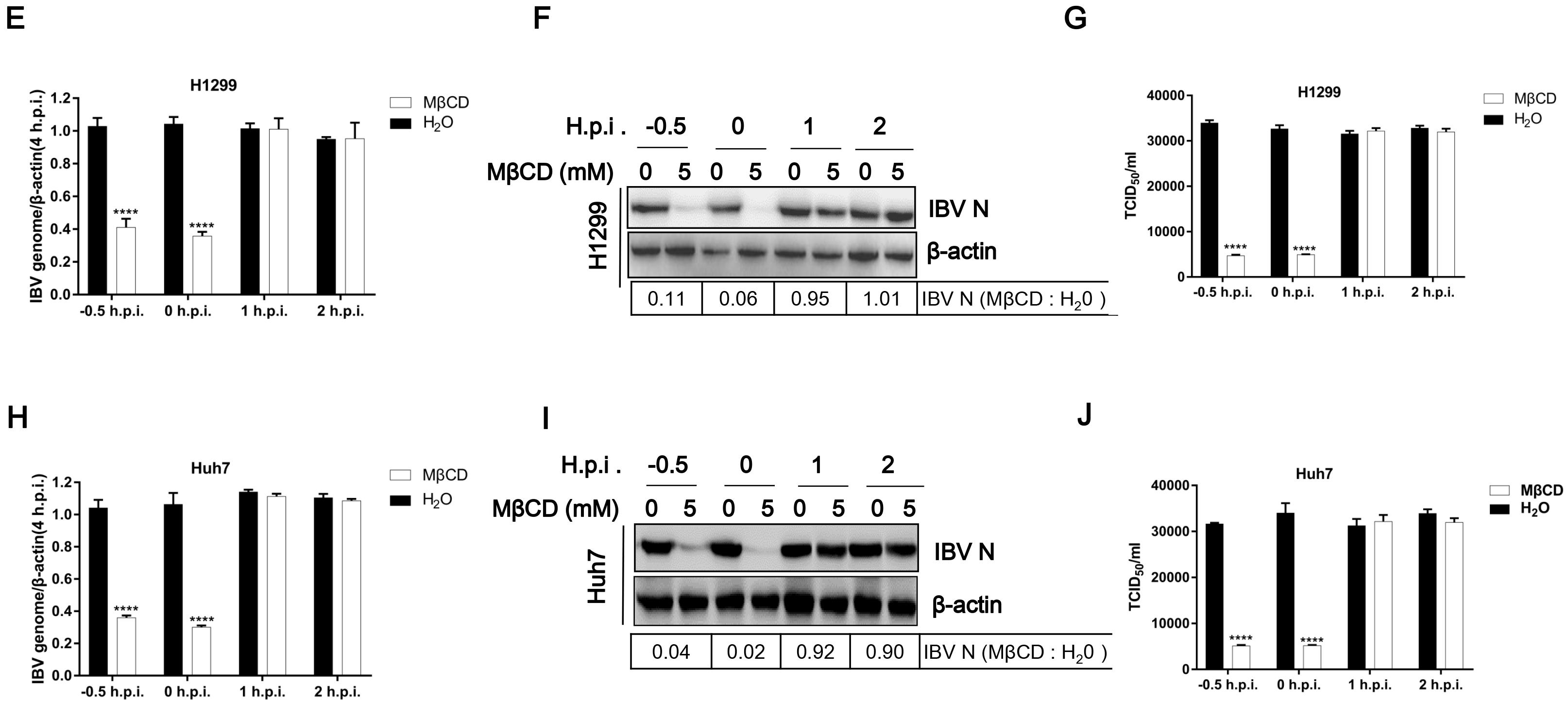
Depletion of lipid rafts before or during virus absorption significantly reduces IBV entry in various cell lines. (A) Time frame of MβCD treatment. (B-I) Vero, H1299, and Huh7 cells were treated with 5 mM MβCD at indicated time points for 30 min, followed with IBV infection. H_2_O treated cells were included in a parallel experiment as control. At 4 h.p.i., the virus genome replication was determined by quantification of viral negative stranded gRNA with semi-quantitative real time RT-PCR (B, E, and H); at 8 h.p.i., the virus N protein expression was determined with Western blot analysis (C, F, and I); at 12 h.p.i. the release of virus particle was determined with TCID_50_ assay (D, Q and J). The virus titers were calculated using the Reed-Muench method. The experiment was performed in triplicate, and average values with stand errors were presented in panel B, D, E, G, H, and J. The intensity of IBV N band was determined with Image J, normalized to β-actin, and shown as fold change of MβCD: H_2_O in table panel of C,F, and I.

To confirm whether the effect of the MβCD on virus entry was due to cholesterol depletion, we examined the effect of exogenous cholesterol replenishment on virus entry after MβCD treatment. Vero, H1299, and Huh7 cells were pretreated with 2.5 or 5 mM MβCD for 30 min, supplemented with/without exogenous 1 mM cholesterol for 1 h, followed with IBV infection. The incoming virus gRNA, the expression of IBV N protein, and the progeny virus particles in the culture medium, were examined. The incoming virus gRNA was examined with semi-quantitative real time RT-PCR at 2 h.p.i.. Results in Fig. 2A, 2D, 2G showed that MβCD treatment reduced virus gRNA internalization, however, cholesterol replenishment restored virus entry. Accordingly, virus protein expression was determined with Western blot at 8 h.p.i.. As shown in Fig. 2B, 2E, and 2H, compared to that in control treatment cells, viral N protein expression was reduced by MβCD treatment and was restored in cholesterol replenishment cells. In accordance, progeny virus release at 12 h.p.i. was rescued by cholesterol replenishment, as determined by TCID_50_ assay (Fig. 2C, 2F, 2I). Above results suggest that IBV attachment and/or entry really depends on intact lipid rafts in various cell lines.

**Figure 2.**
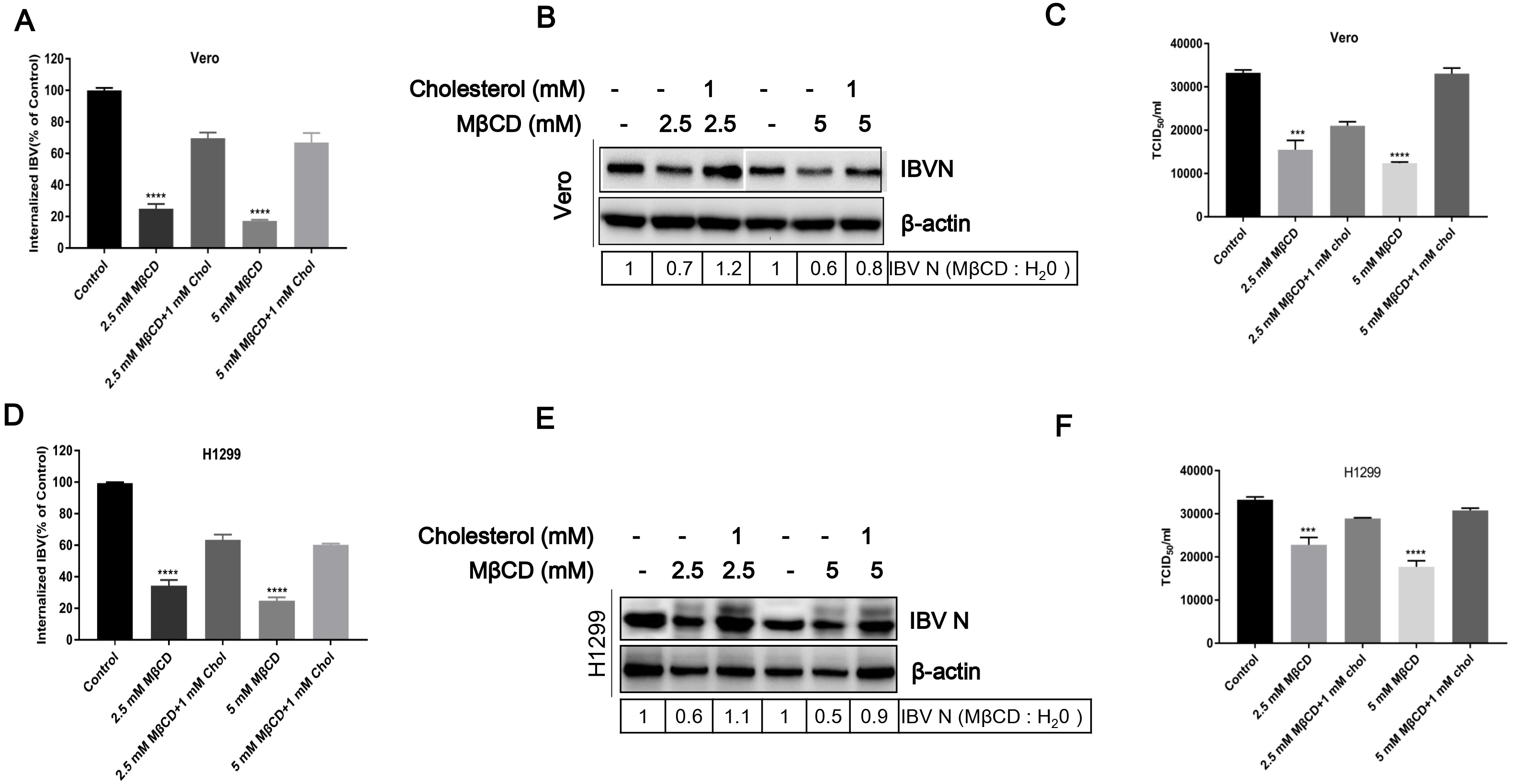
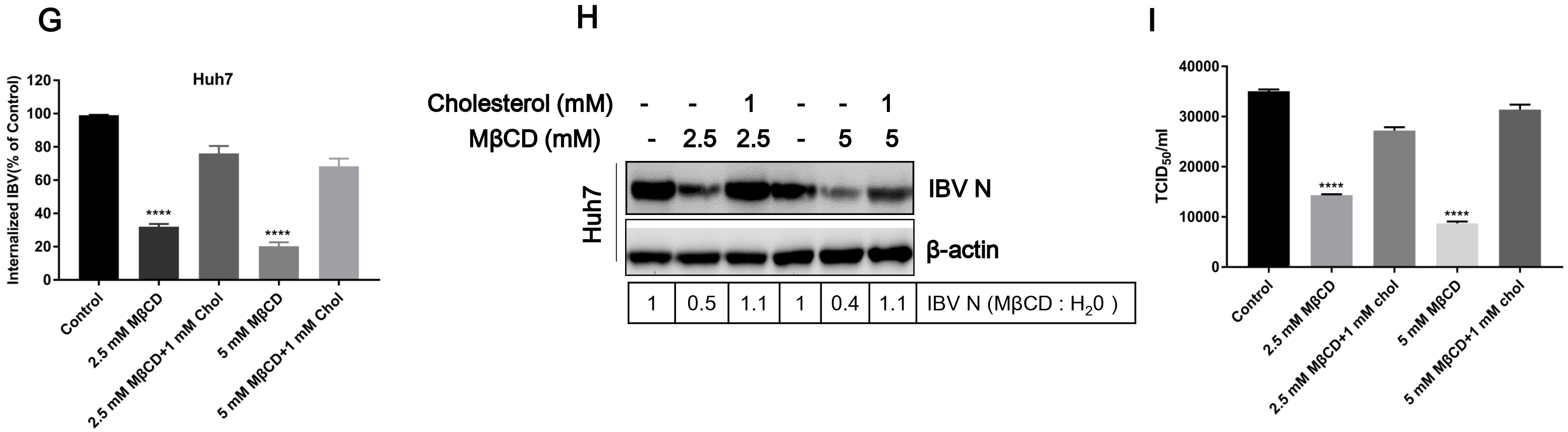
Exogenous cholesterol replenishment restores virus infection. Vero, H1299 cells, and Huh7 cells were pretreated with 2.5 mM or 5 mM MβCD and supplemented with 1 mM cholesterol before IBV infection. Mock-treated and MβCD-treated cells without cholesterol replenishment were included in a parallel experiment as control. The entry of IBV was measured by quantification of viral gRNA at 2 h.p.i. (A, D, and G); the expression of IBV N protein was measured at 8 h.p.i. (B, E, and H); the release of virus particles in the medium was measured at 12 h.p.i., (C, F, and I). The virus titer was calculated using the Reed-Muench method. The experiment was performed in triplicate, and average values with stand errors were presented in panel A, C, D, F, Q and I. The intensity of IBV N band was determined with Image J, normalized to β-actin, and shown as fold change of MβCD: H_2_O in table panel of B, E, and H..

### Association of IBV with lipid rafts during the early stage of infection

It should be mentioned that the receptor of IBV entry is controversial. Whereas some work states that heparan sulfate is a selective attachment factor for the IBV Beaudette strain (48), another report concludes that sialic acid is a receptor determinant of infection (49). Since membrane cholesterol plays an important role in early infection stages, the interaction of IBV with its receptor may occur in lipid rafts. To test this hypothesis, Vero cells were incubated with IBV for 1 h at 4°C and lipid rafts were isolated by a sucrose flotation gradient. The individual fractions were analyzed for the presence of rafts marker flotillin-1, IBV N protein, and IBV S protein by Western blotting. In IBV-infected cells, we found that S protein and N protein were co-localized with the rafts marker flotillin-1in fractions 2 and 3 (Fig. 3A). Thus, the biochemical fractionation analysis suggests the association of IBV with lipid rafts during attachment.

**Figure 3.**
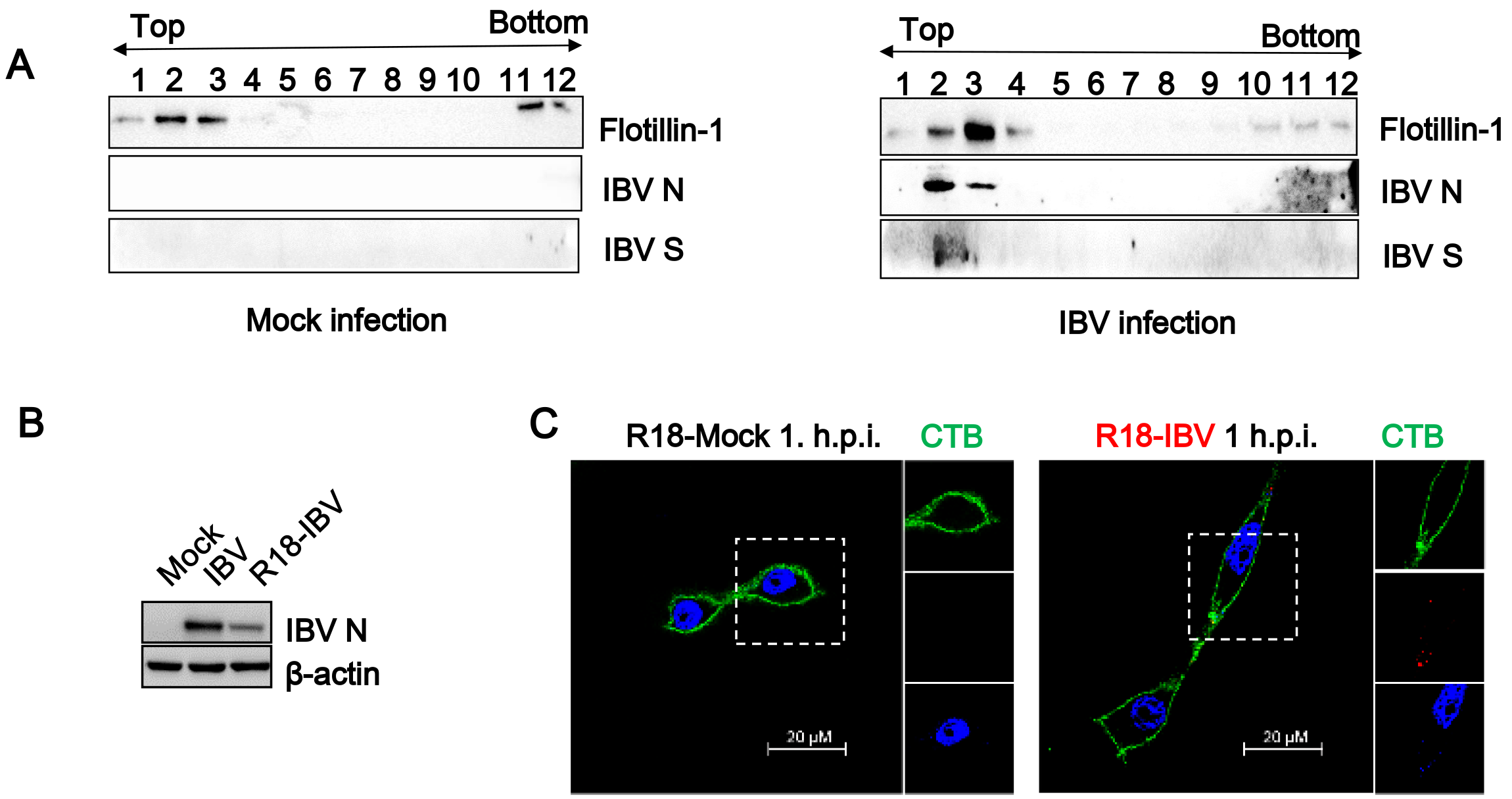
IBV associates with lipid rafts during attachment. (A) Vero cells were incubated with IBV (MOI 5) at 4°C for 1 h, followed by treatment with TNE buffer with 1% TX-100 at 4°C for 30 min. Cells were subjected to membrane flotation analysis. Mock infection was included in a parallel experiment as control. The isolated membrane fractions were collected from top to bottom, and lipid rafts maker Flotillin-1, IBV N protein and S protein were detected by Western Blot analysis. (B) Vero cells were infected with 5 MOI of IBV or R18-IBV and the expression of IBV N protein was analyzed with Western blot at 8 h.p.i.. (C) Vero cells were incubated with 5 MOI of R18-IBV (red) and Alex Fluor 488 conjugated CTB (bind to lipid rafts marker GM1, green) for 1 h at 4°C, followed with DAPI staining (blue) for 15 min. Images were acquired by LSM880 confocal laser-scanning microscope (Zeiss). Representative images were shown. Scale bars=20 μm.

To directly observe the binding of virus particles to cell surface lipid rafts, IBV particles were concentrated and labeled with the lipophilic fluorescent dye R18. The cell culture supernatant was concentrated and mixed with R18 as mock group. Such R18 labeled virus retained high levels of infectivity (Fig. 3B). Immunofluoresence was then performed to check the interaction between R18-IBV and lipid rafts marker GM1. Vero cells were incubated with either mock or R18-IBV, and CTB-FITC (binds to lipid rafts marker GM1) at 4°C for 1 h, followed with DAPI staining of nucleus. As shown in Fig. 3C, the mock infected cells displayed no R18 red signal, while the R18-IBV infected cells displayed red dot signal on the plasma membrane, and co-localized with CTB-FITC, which represents the outline of lipid rafts. This result confirms the association of IBV with lipid rafts on cell surface.

### IBV entry is low pH dependent

Although previous study demonstrates that IBV strain Beaudette entry is sensitive to drug treatment that neutralize the low pH of endosomes in BHK cells (44), it has also been shown that different cell types have different abilities to allow viral entry (50). Thus, to dissect the IBV entry pathway, it is necessary to verify whether the low pH dependent entry of IBV is cell type specific. To this end, we increased the pH in Vero, BHK, H1299, and Huh7 cells by using endosome acidification inhibitor NH_4_Cl, and checked IBV internalization/replication. To ensure subtoxic doses of the drug, Vero cells were treated with 0-160 mM NH_4_Cl for 30 min, and cytotoxic effects were evaluated by cell viability assay. As shown in Fig. 4A, Vero cells tolerated the treatment of NH_4_Cl up to 100 mM. Next, acridine orange staining was performed to confirm increasing pH in intercellular vesicles caused by the drug treatment. Results in Fig. 4B showed that untreated cells displayed typical orange fluorescence, indicative of acidic compartments. As expected, the orange fluorescence decreased along with increasing concentration of NH_4_Cl treatment, and 100 mM NH_4_Cl treatment totally quenched the orange fluorescence, confirming pH increased in NH_4_Cl treated cells. To examine the functional role of a low pH on IBV infectivity, we pretreated various cell lines with 0-100 mM NH_4_Cl for 30 min, followed by inoculation with IBV for 1 h at 4°C. The temperature was shifted to 37 °C for an additional 1 h to allow synchronized entry. Internalized virus was quantified by detection of viral gRNA at 2 h.p.i.. As shown in Fig. 4C, 4E, 4G; 4I, virus internalization was significantly reduced by NH4Cl pretreatment in a dose-dependent manner, compared to those in H_2_O treated cells. The virus replication upon NH_4_Cl treatment was monitored by detection of viral N protein at 8 h.p.i.. The basic mechanism of pH-dependent endocytosis for VSV has been known for many years and is well documented (51-54). Thus, VSV infection of Vero cells was included in this experiment as a positive control. Western blot result in Fig. 4D showed that VSV G protein expression was decreased by NH4Cl treatment in dose dependent manner, suggesting that this drug really inhibited VSV infection. Data in Fig. 4D, 4F, 4H, 4J showed that IBV N protein production was significantly decreased by NH_4_Cl treatment in dose dependent manner. Taken together, these results demonstrate that IBV entry requires low-pH environment in the endosome in various cell lines, consistent with the previous report (44).

**Figure 4.**
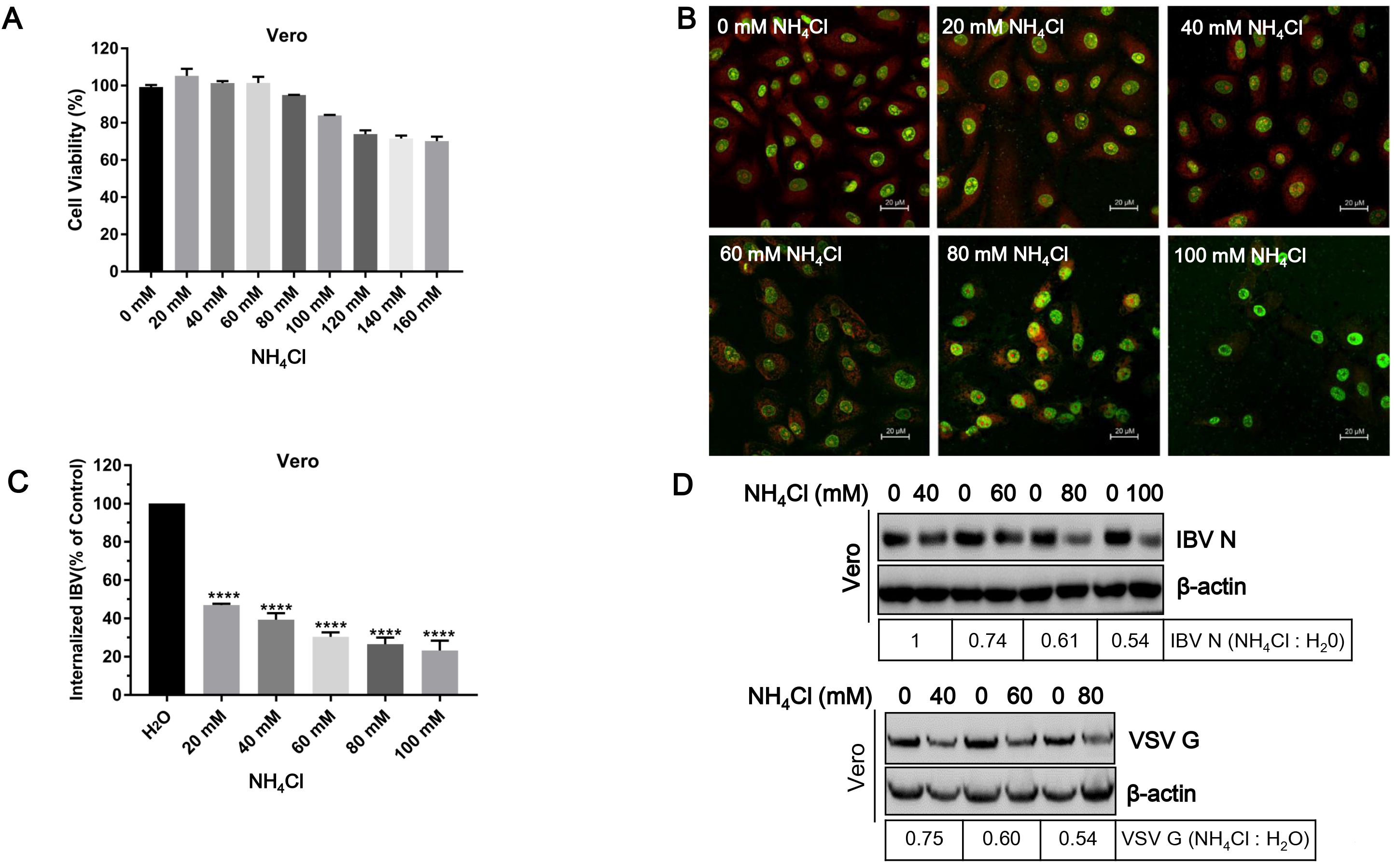
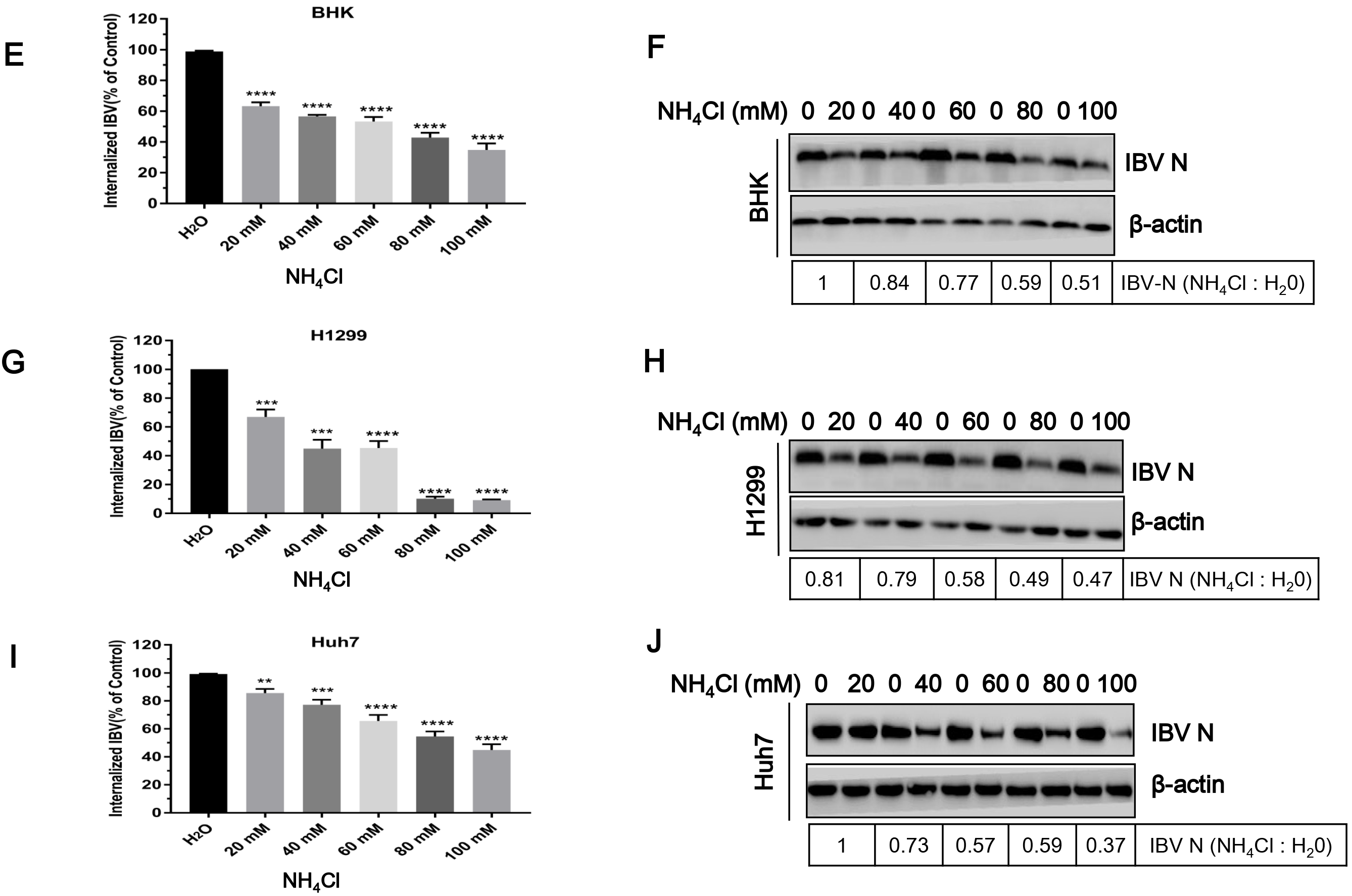
IBV entry is hampered by increase of pH in the intracellular vesicles. (A) Vero cells were treated with 0-160 mM NH_4_Cl for 30 min, and the cell viability was assessed with WST-1 cell proliferation and cytotoxicity assay kit. (B) Cells treated with NH_4_Cl (0-100 mM) were stained with acridine orange and DAPI, and observed under fluorescence microscope. Representative images were shown. Scale bars=20 μm. (C-J) Vero, BHK, H1299, and Huh7 cells were treated with increasing concentrations (0-100 mM) of NH_4_Cl for 30 min, followed with IBV infection (1 MOI). The incoming virus level was determined by quantification of viral gRNA at 12. p.i. (C, E, G; and I); virus N protein expression was determined by Western blot analysis at 8 h.p.i. (D, F, H, and J). Vero cells pretreated with NH_4_Cl followed with VSV infection were set as control. The expression of VSV G protein was detected with Western blot analysis (lower panel in D). The experiment was performed in triplicate, and average values with stand errors were presented in panel A, C, E, G, and I. The intensity of IBV N or VSV G band was determined with Image J, normalized to β-actin, and shown as fold change of NH_4_Cl: H_2_O in table panel of D, F, H, and J.

### IBV enters cells via clathrin-medicated endocytosis (CME)

Several pH-dependent endocytic pathways are used by various viruses, including CME, CavME, macropinocytosis, and clathrin/caveolin-independent endocytosis (55). To identify the endocytic pathway employed by IBV, pharmacological inhibition, siRNA-mediated depletion, and overexpression of dominant negative or constitutive active proteins were carried out to block or strengthen respective endocytic pathway. First, we investigated the effect on IBV entry by diverse pharmacological drugs treatments which have been well known to inhibit CME, CavME, or macropinocytosis. Vero, H1299, and DF-1 cells were exposed to increasing concentrations of Chlorpromazine (CPZ), an inhibitor of clathrin lattice polymerization (56), Amiloride, a specific inhibitor of Na^+^/H^+^ exchanger activity important for macropinosome formation(17, 23, 57, 58), or Nystatin, a CavME inhibitor (59). Virus endocytosis was quantified by measuring the amount of intracellular viral gRNA at 2 h.p.i., and virus replication was scored by IBV N protein expression level at 8 h.p.i.. The toxicity of each inhibitor was carefully determined (data not shown). As semi-quantitative real time RT-PCR results shown in Fig. 5A, 5C, and 5E, compared to the control group, pretreatment of cells with CME inhibitor CPZ strongly inhibited the internalization of viral gRNA in Vero, H1299, and DF-1 cells, which is dose dependent; however, Amiloride pretreatment did not significantly change the internalization of virus in all three cell lines. Nystatin pretreatment did not alter the virus internalization in either Vero or DF-1 cells; however, it slightly reduced the internalization in H1299 cells. Western blot detection of viral N protein expression in Fig. 5B, 5D, and 5F showed that CPZ significantly inhibited the viral N protein expression in a dose dependent manner; Amiloride pretreatment did not significantly change the expression of N protein in all three cell lines. In Nystatin pretreated Vero cells, IBV N protein remained in a steady level, compared to control treated cells; however, in H1299 and DF-1 cells, Nystatin pretreatment slightly reduced the viral protein expression. It has been reported that VSV entry is mediated by CME, and indeed, CPZ pretreatment of Vero cells reduced the VSV G protein expression in a dose dependent manner, while Amiloride and Nystatin pretreatment had no effect (Fig. 5I). Thus, these results reveal that IBV enters cells via CME, but not via macropinocytosis. Whether the CavME is involved in IBV endocytosis may be cell type dependent and is controversial. To explore the involvement of CavME in IBV entry, we utilized Huh7 cell, which lacks functional caveolae (15, 60), to perform above experiment. As evidenced by high level of intracellular viral gRNA and N protein expression, efficient virus internalization and replication was observed in Huh7 cells (Fig. 5G-H), demonstrating that IBV entry is independent of caveolae. Moreover, CPZ pretreatment significantly reduced IBV gRNA internalization and N protein expression in Huh7 cells, while neither Amiloride nor Nystatin reduced the internalization and replication (Fig. 5G-H), consistent with those observed in other cell lines. Combining above results, we conclude that IBV entry is mediated by CME, and this is not restricted to cell types.

**Figure 5.**
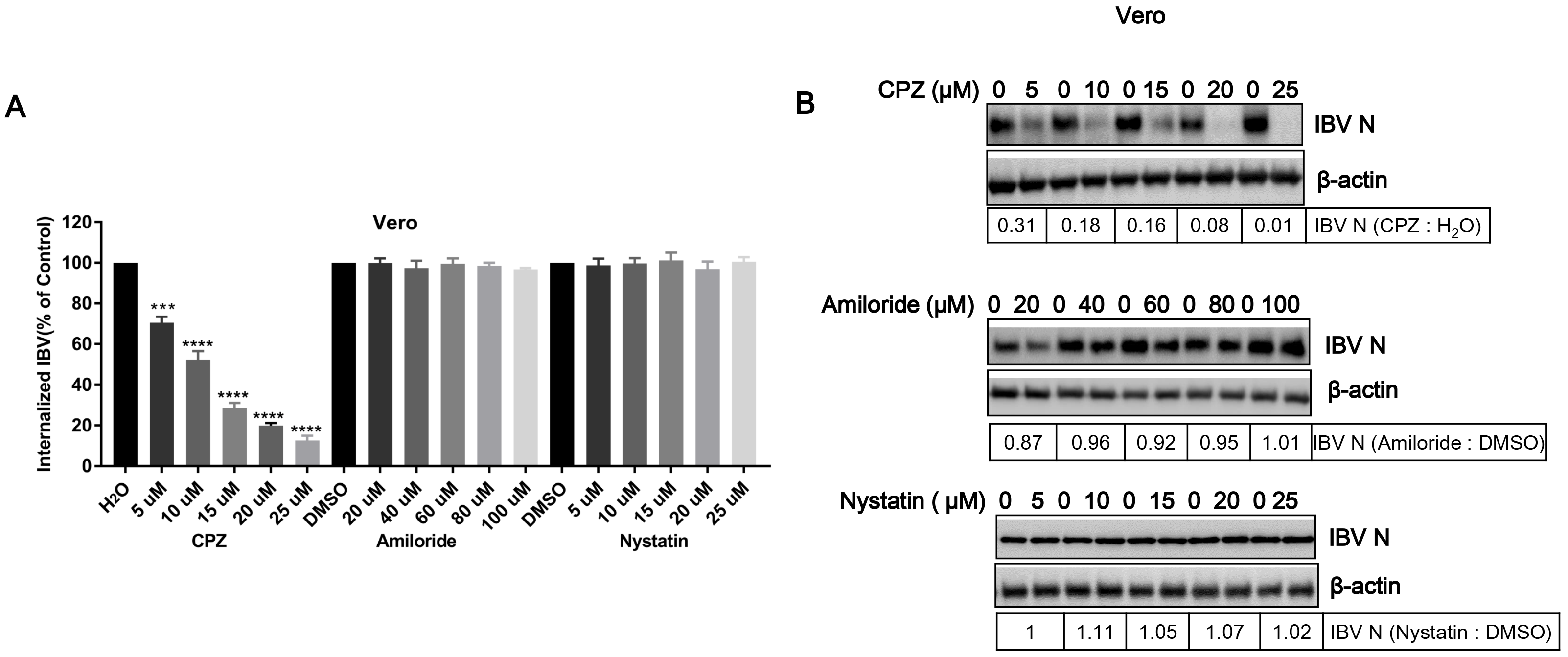
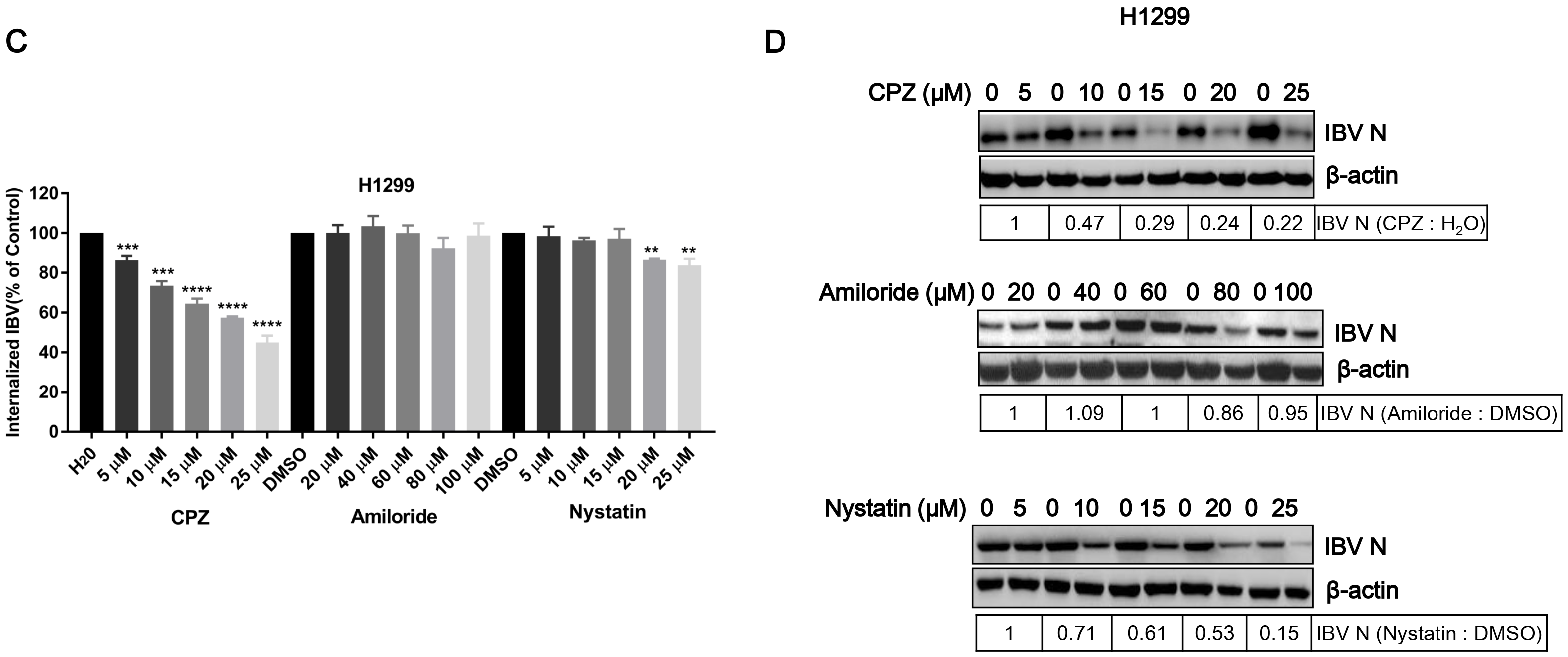
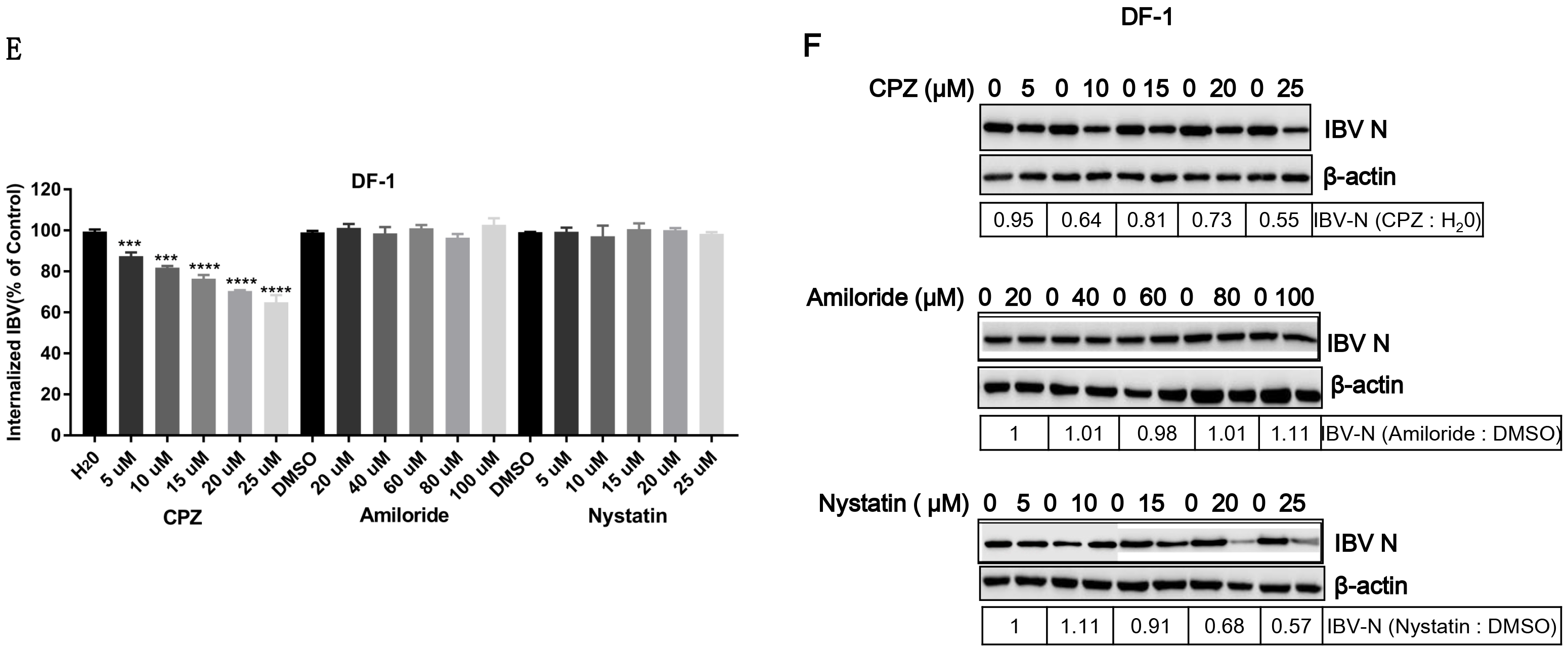
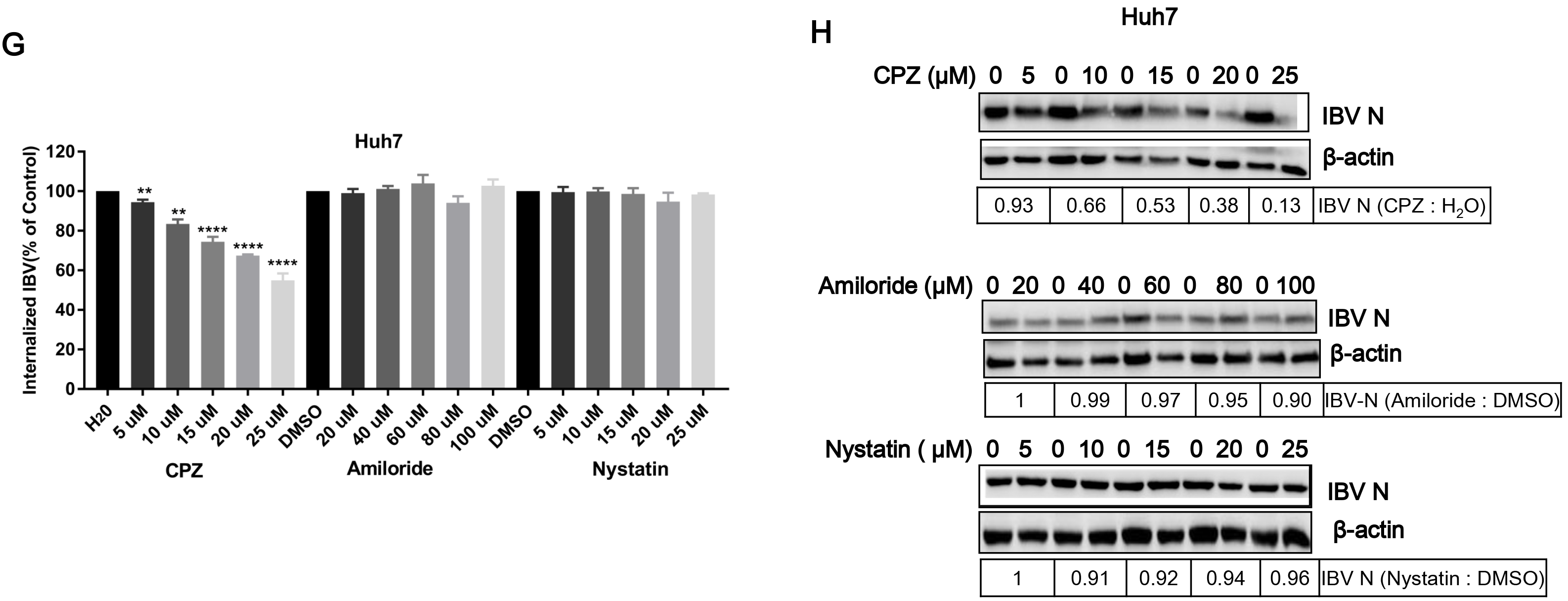
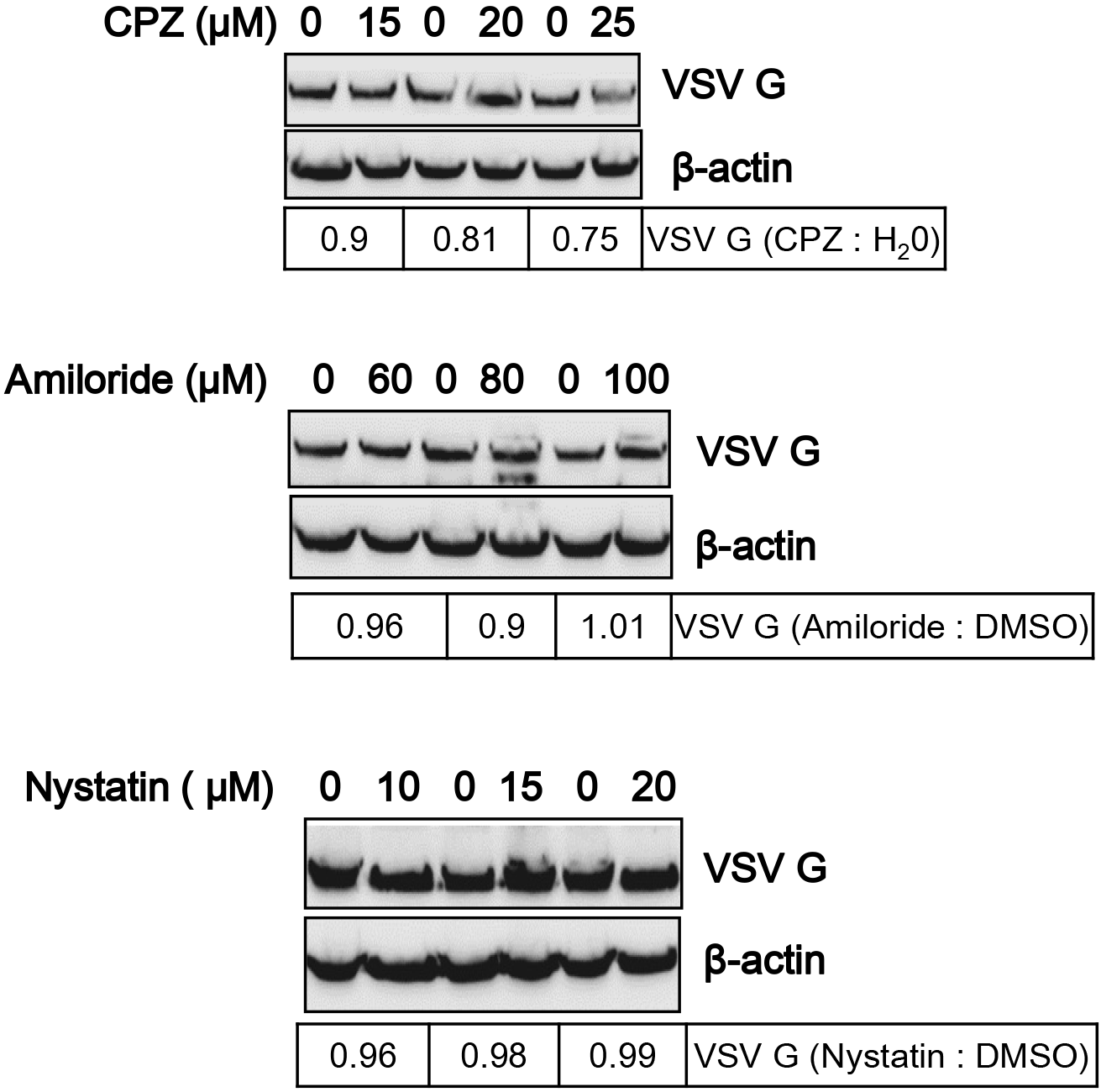
IBV entry is blocked by clathrin-mediated endocytosis (CME) inhibitor CPZ. (A-H) Vero, H1299, DF-1, and Huh7 cells were pretreated with increasing concentrations of CPZ (CME inhibitor), Amiloride (macropinocytosis inhibitor), or Nystatin (CavME inhibitor) for 30 min and infected with 1 MOI of IBV. H_2_O or DMSO pretreatment in a parallel experiment was set as control. The incoming virus level was determined by quantification of viral gRNA at 2 h.p.i. (A, C, E, and G); the expression level of IBV N protein was analyzed at 8 h.p.i. (B, D, F, and H). The experiment was performed in triplicate, and average values with stand errors were presented in panel A, C, E, and G. (I) Vero cells pretreated with drugs and infected with 1 MOI of VSV were set as control. The expression of VSV G protein was analyzed by Western blot at 8 h.p.i.. The intensity of IBV N or VSV G band was determined with Image J, normalized to β-actin, and shown as fold change of CPZ: H_2_O, Amiloride: DMSO, or Nystatin: DMSO in table panel of B, D, F, H, and I.

### CME of IBV is dependent on dynamin1 but not Eps15

Eps15 is a regulatory protein which has been characterized by a ubiquitous and constitutive association with the AP-2 protein adapter. Microinjection of Eps15 antibody interfered with transferring and EGFR internalization (61), suggesting that Eps15 plays an important role in CME. Several reviews described previously that the expression of the C-terminal dominant negative form of Eps15-DN, which contains the AP2-binding site, could efficiently block the uptake of Sindbis virus (62). Dynamin1 is a GTPase facilitating membrane fission to generate endocytic vesicles in CME and CavME (47, 63, 64). Many reviews describe that dynamin1 is required for the internalization of various viruses, as the dynamin1-K44A, a dominant negative mutant demonstrated preciously to block virus entry (65, 66). To examine the involvement of Eps15 and dynamin1 in IBV entry, we expressed pEGFPN1, GFP-tagged Eps15-WT, Eps15-DN, dynamin1-WT, and dynamin1-DN in H1299 cells, respectively, followed by IBV infection. pEGFPN1 was expressed as the control. Vero cells were not chosen for the protein expression experiment due to their low transfection efficiency. The successful expression of GFP, GFP-tagged Eps15-WT, GFP-Eps15-DN, dynamin1-WT, and dynamin1-DN was detected in Fig. 6A (low panel), and the expression level of Eps15 is lower than dynamin1. Compared to that in GFP expressing cells, the expression of IBV N protein was increased in dynamin1-WT expressing cell, whereas reduced in dynamin1-DN expressing cells, demonstrating the crucial role of dynamin1 in IBV infection. Surprisingly, neither Eps15-WT nor Eps15-DN changed the expression level of IBV N, due to unknown reason. It has been reported that VSV entry relies on functional Eps15 (67), thus, the effect of Eps15 on VSV entry of Vero cells was also examined as control experiment. Eps15-DN indeed inhibited VSV infectivity, as evidenced by the reduced level of VSV G protein (by 0.66-fold) in Eps15-DN expressing cells (Fig. 6D). Therefore, the failure of blocking IBV infectivity by Eps15-DN is not due to low expression level of Eps15-DN or experimental error. To confirm above result, the internalization of IBV gRNA was quantified at 2 h.p.i.. As shown in Fig. 6B, expression of dynamin1-DN significantly blocked virus internalization by 0.6-fold, whereas expression of dynamin1-WT, Eps15-WT, and Eps15-DN had no significant effect on virus entry. The release of progeny virus particle was also determined by TCID_50_. Results in Fig. 6C showed that expression of dynamin1-DN significantly reduced progeny virus production by around 0.5-fold, but dynamin1-WT, Eps15-WT, and Eps15-DN had no significant effect on this, compared to those in GFP expressing cells. In all, above results suggest that IBV entry depends on functional dynamin1, but not Eps15.

**Figure 6.**
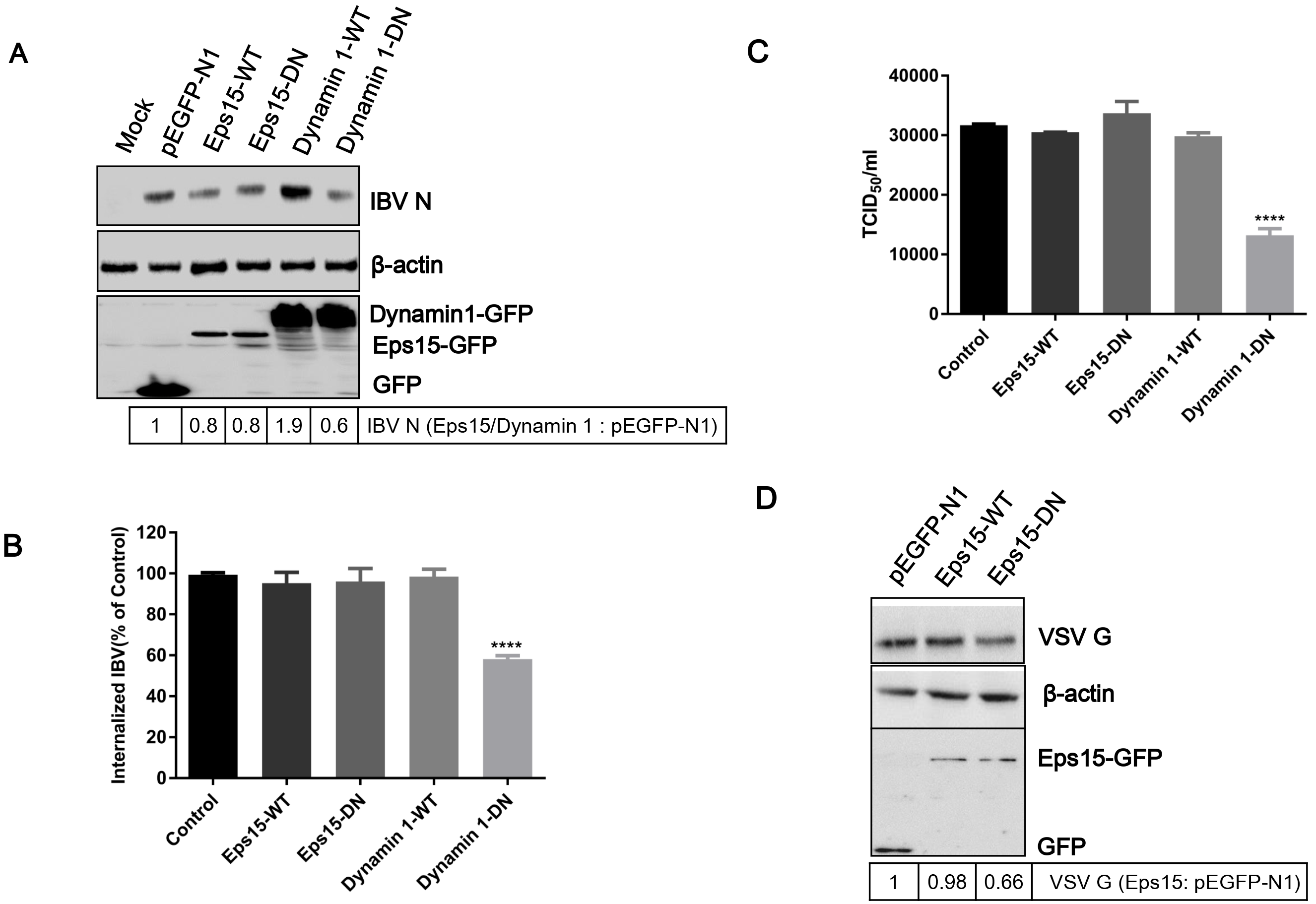
IBV endocytosis is dependent on dynamin1. (A-C) H1299 cells were transfected with constructs encoding GFP fused Eps15-WT, Eps15-DN, Dyanmin1-WT, Dynamin1-DN, or vector pEGFP-N1, respectively. At 24 h post-transfection, cells were infected with IBV (1 MOI). Cells were harvested at 8 h.p.i.. The expression of GFP, GFP-Eps15-WT, GFP-Eps15-DN, GFP-Dynamin1-WT, GFP-Dynamin1-DN, and IBV N protein was checked with Western blot analysis by using anti-GFP or anti-IBV N (A); the internalized IBV genome was analyzed with real time RT-PCR at 2 h.p.i. (B); the virus particle release was determined by TCID_50_ at 12 h.p.i. (C). The experiment was performed in triplicate, and average values with stand errors were presented in panel B and C. (D) Vero cells transfected with constructs encoding GFP fused Eps15-WT, Eps15-DN, or vector pEGFP-N1, respectively. At 24 h post-transfection, cells were infected with VSV (1 MOI). Cells were harvested at 8 h.p.i.. The expression of GFP, GFP-Eps15-WT, GFP-Eps15-DN, and VSV G protein was checked with Western blot analysis by using anti-GFP or anti-VSV G. The intensity of IBV N or VSV G band was determined with Image J, normalized to β-actin, and shown as fold change of Eps15/Dynamin 1: pEGFP-N1 or Eps15: pEGFP-N1 in table panel of A and D.

To further confirm above conclusion, we analyzed the effect of Eps15 and dynamin1 depletion on IBV uptake. To this purpose, cells were transfected with siRNA targeting to either Eps15 or dynamin1, and then infected with IBV (H1299 cells) or us VSV (Vero cells). The knock down efficiency of Eps15 or dynamin1 and intracellular IBV gRNA were measured by semi-quantitative real time RT-PCR. As shown in Fig. 7, successful silence of Eps15 mRNA was obtained by siEps15 in both H1299 and Vero cells (Fig. 7A, C); however, silence of Eps15 expression had no significant effect on the IBV gRNA internalization (Fig. 7B). Moreover, although Eps15 depletion inhibited VSV G expression (Fig. 7D), IBV N protein expression was unchanged by this knock down (Fig. 7C). These data further confirmed although IBV entry is dependent on CME, Eps15 is not required. Furthermore, knock down of dynamin1 (by 92%) was accompanied by a significant reduction of the amount of incoming IBV gRNA (by 48%) and IBV protein expression (by 71%) (Fig. 7E-G). Collectively, these results indicate that dynamin1, but not Eps15, is functionally required for IBV entry.

**Figure 7.**
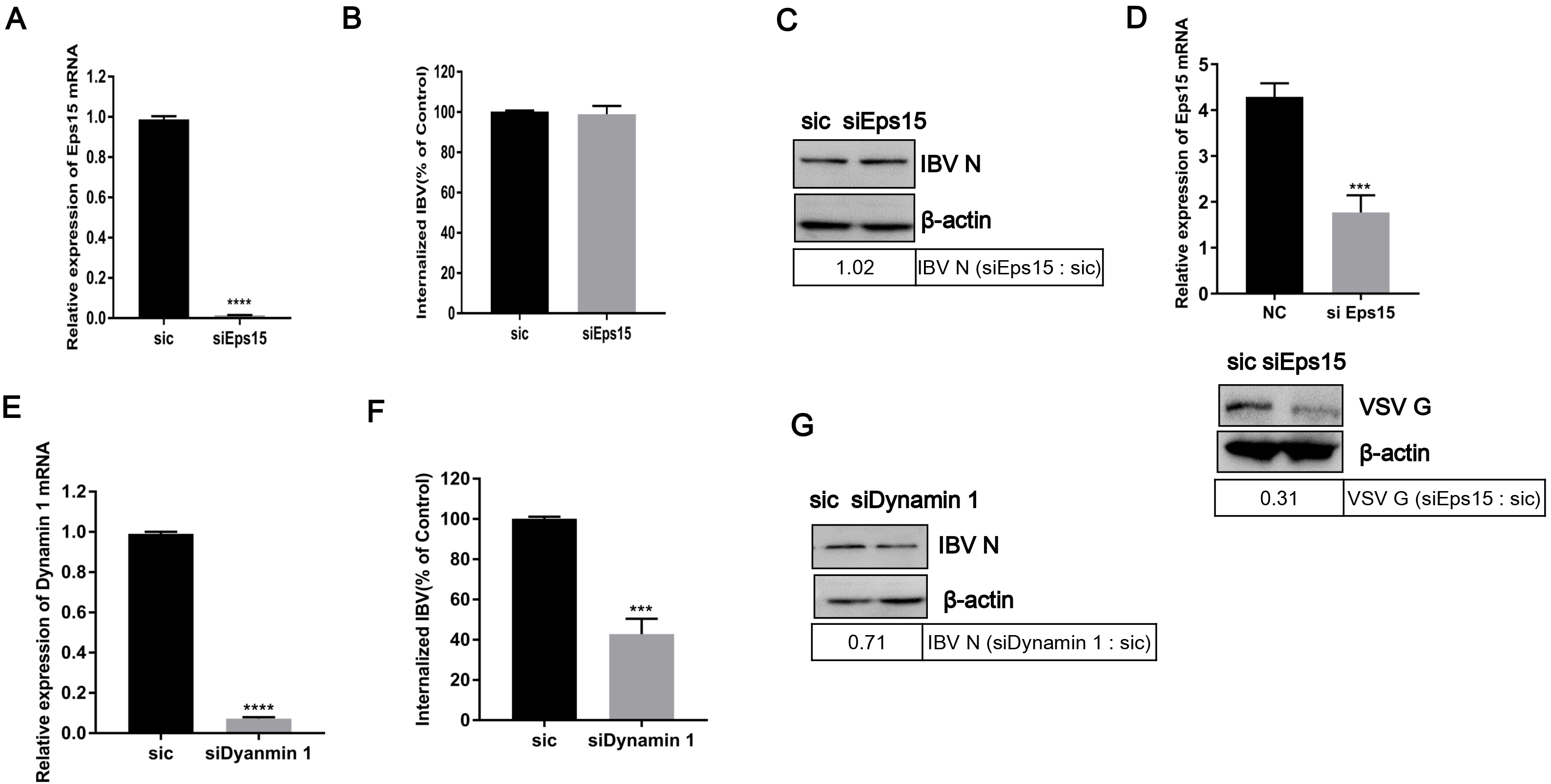
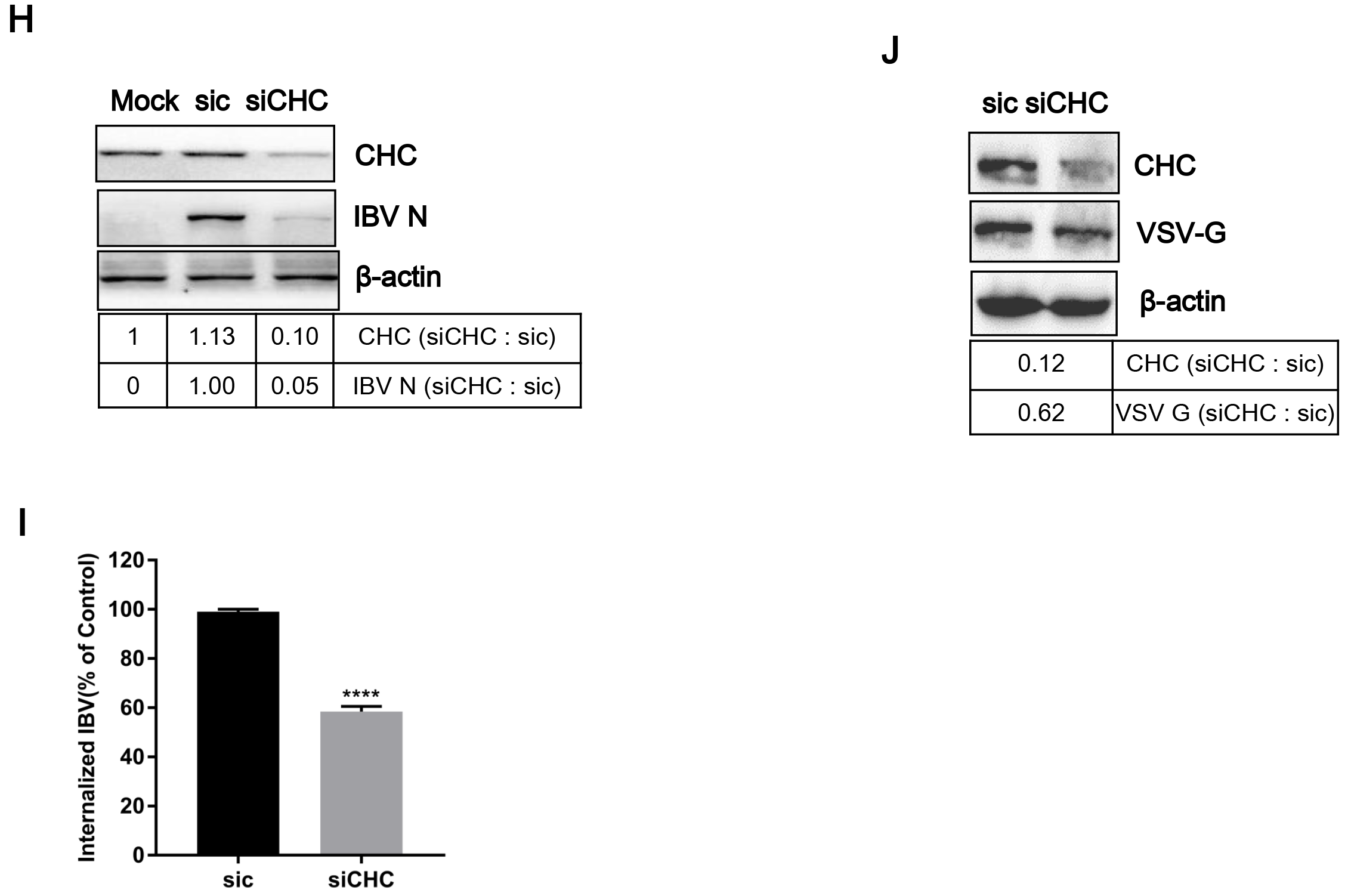
Both dynamin1 and CHC are involved in IBV endocytosis. (A-D) Cells were transfected with non-target siRNA (sic) or Eps15 siRNA (siEps15) for 36 h, followed with IBV infection (H1299 cells) or VSV infection (Vero cells). The knock down effect of Eps15 and the internalization of IBV was analyzed by semi-quantitative real time RT-PCR at 2 h.p.i. (A, B, and upper panel in D); the IBV N protein or VSV G protein expression was determined with Western blot at 8 h.p.i. (C and lower panel in D). (E-G) H1299 cells were transfected with non-target siRNA (sic) or Dynamin1 siRNA (siDynamin1) for 36 h, followed with IBV infection. The knock down effect of Dynamin1 and the internalization of IBV gRNA were analyzed by semi-quantitative real time RT-PCR at 2 h.p.i. (E and F); the expression level of IBV N protein was determined with Western blot at 8 h.p.i.(G). (H-J) Cells were transfected with siCHC for 36 h, followed by IBV infection (H1299 cells) or VSV infection (Vero cells). The expression level of CHC, IBV N protein, or VSV G protein was detected by Western blot at 8 h.p.i. (H and J); IBV internalization was analyzed at 2 h.p.i. (I). The experiment was performed in triplicate, and average values with stand errors were presented in panel A, B, E, F and I. The intensity of IBV N, VSV G; or CHC band was determined with Image J, normalized to β-actin, and shown as fold change of siEps15: sic, siDynamin1: sic, or siCHC: sic in table panel of C, D, G; H, and J.

We further analyzed the effect of silencing CHC expression on IBV uptake and infection. To this purpose, cells were transfected with siRNA targeting CHC and then infected with IBV (H1299 cells) or VSV (Vero cells). As shown in Fig. 7H-J, depletion of CHC was accompanied by a significant reduction of IBV gRNA internalization and IBV/VSV protein expression, which further demonstrates a functional clathrin pathway is required for efficient IBV infection.

### IBV entry leads to active actin rearrangement

In *Saccharomyces cerevisae*, clathrin-coated patches on the cell surface mature into ~200 nm tubular invaginations with coat proteins at their tip. The actin cytoskeleton machinery is recruited after coat formation and provides an essential force for membrane deformation and internalization (69). Recent studies have revealed that the rhabdoviruses (70), VSV (7), RABV (71), and IHNV (72) are internalized through CME and depend on actin. Although we have demonstrated that IBV enter cells by hijacking clathrin dependent route, it is unclear whether actin filaments are involved in IBV uptake. To examine the role of actin filaments in IBV entry, we first examined the actin polymerization during IBV internalization. Vero cells were incubated with IBV at 4°C for 1 h, and the temperature was shifted to 37°C. Actin filaments were stained with Alexa Fluor 488-phalloidin (green) and IBV N antibody at 15 min time interval and were observed by confocal microscopy. In mock-infected cells, the actin stress fibers were visible clearly. At 15 to 45 min.p.i., the number of actin stress fibers dramatically decreased, the cells rounded up, and cells surface displayed significant blebbing. The cell morphology and actin distribution returned to normal at 60 min.p.i. (Fig. 8A). Above observation indicates that IBV infection results in transient actin de-polymerization and re-arrangement. IBV N protein antibody is not sensitive enough to detect the incoming virus. To check whether of IBV internalization associates with actin, R18-IBV was applied to Vero cell. Actin filaments were stained with Alexa Fluor 488-phalloidin at 15 min time interval, and the red signal of R18-IBV and green signal of actin were observed. As expected, images in Fig. 8B showed that there was no R18 red signal in mock infection group. In the R18-IBV infection group, R18-IBV was detected at cell surface at 0 min, displayed as red signal. After attachment, R18-IBV entered cells and co-localized well with actin at 15, 30, and 45 min, shown as yellow signal. At 60 min, Most R18-IBV particles separated with actin, displayed as red signal again. This observation suggests that IBV enters cells via actin cytoskeleton. Here, we also incubated the well characterized R18-VSV with Vero cells as control group. As R18-VSV enters into cells quickly, we chose the time interval of 5 min. Image in Supplementary Fig. 2A showed that R18-VSV also caused actin fiber de-polymerization and cell rounded up at 5-30 min.p.i.. The cell morphology and actin distribution returned to normal at 30 min. Meanwhile, R18-VSV co-localized well with actin at 5, 10, 20, and 30 min, suggesting that VSV also moves along actin fiber after internalization, consistent with previous report (73).

**Figure 8.**
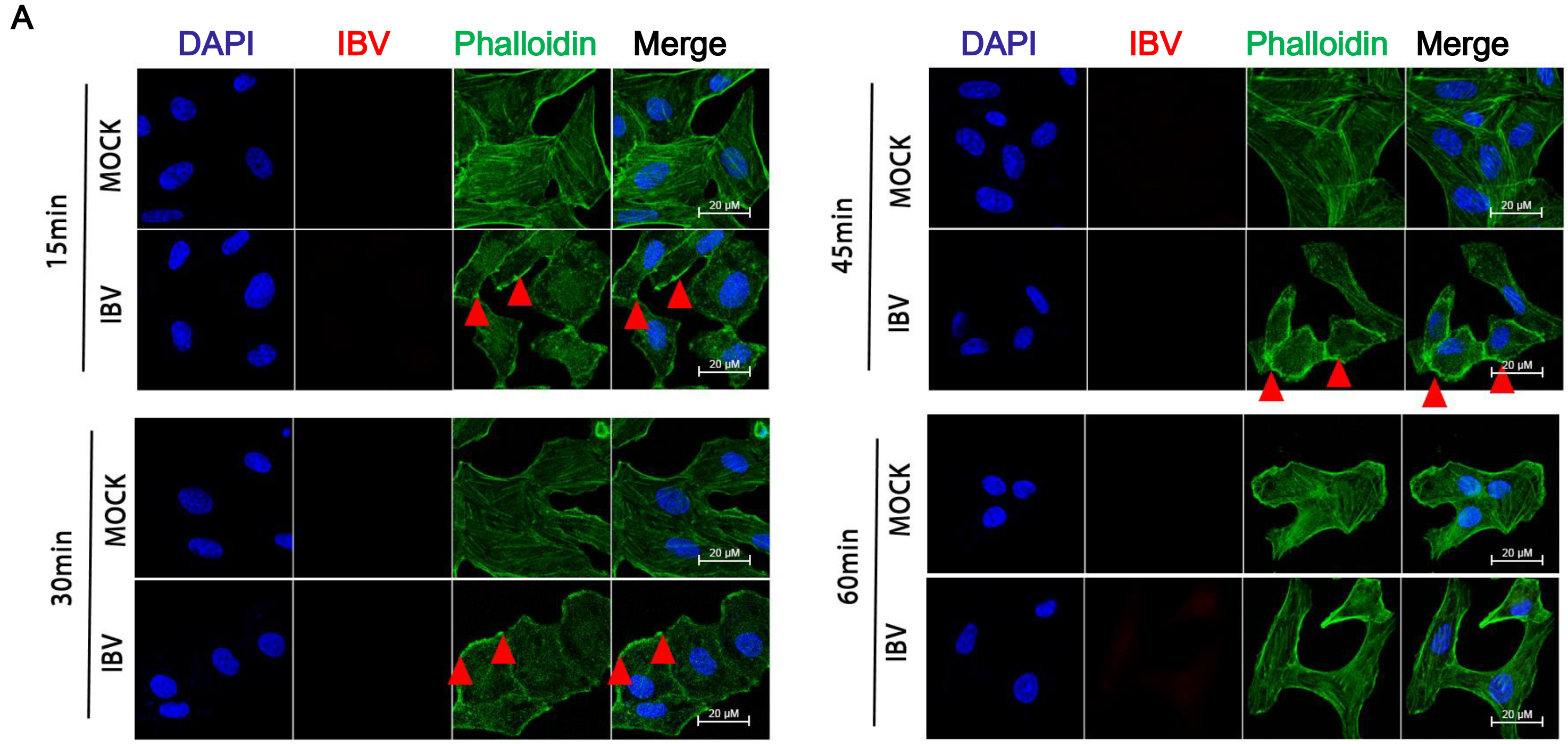
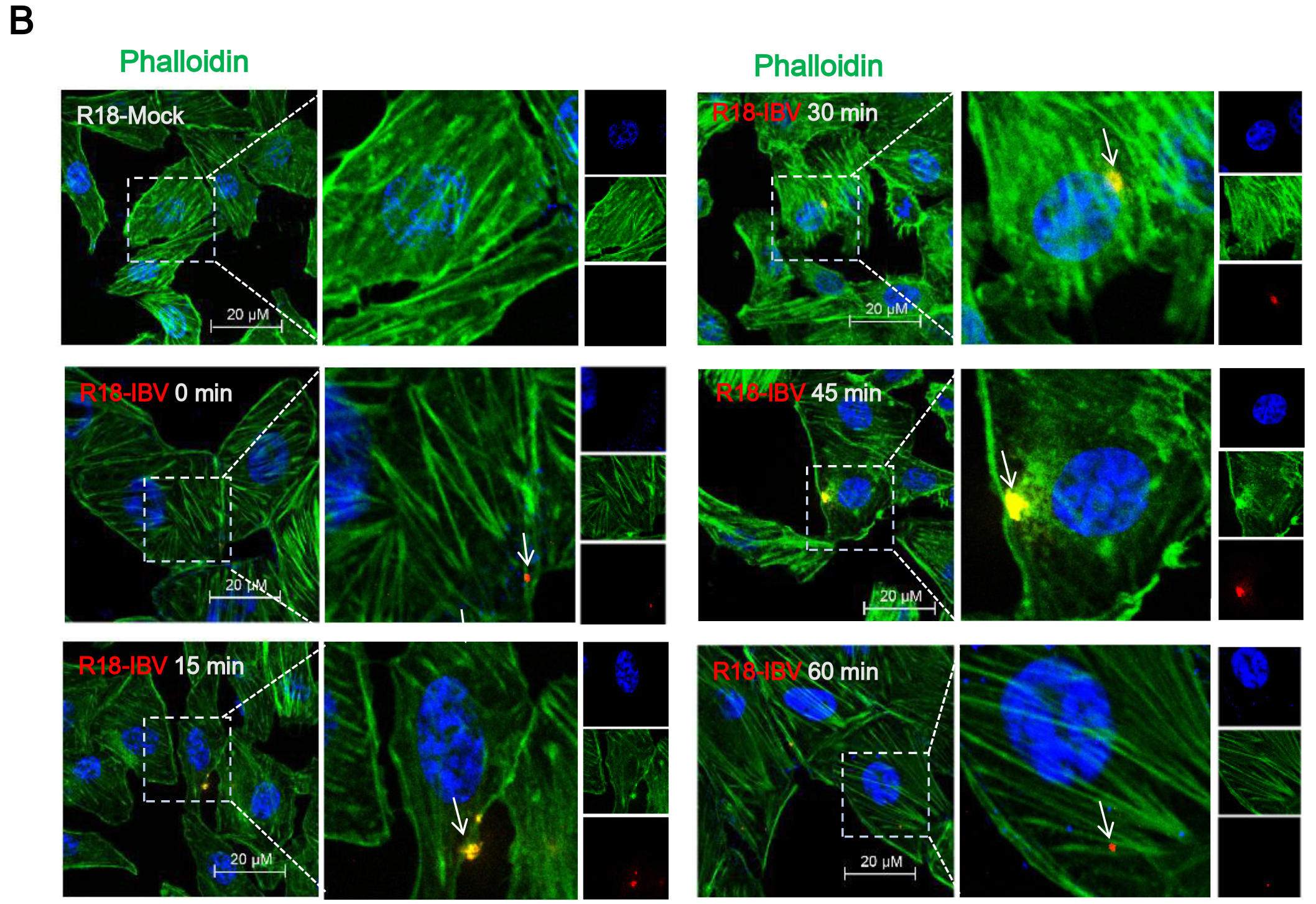
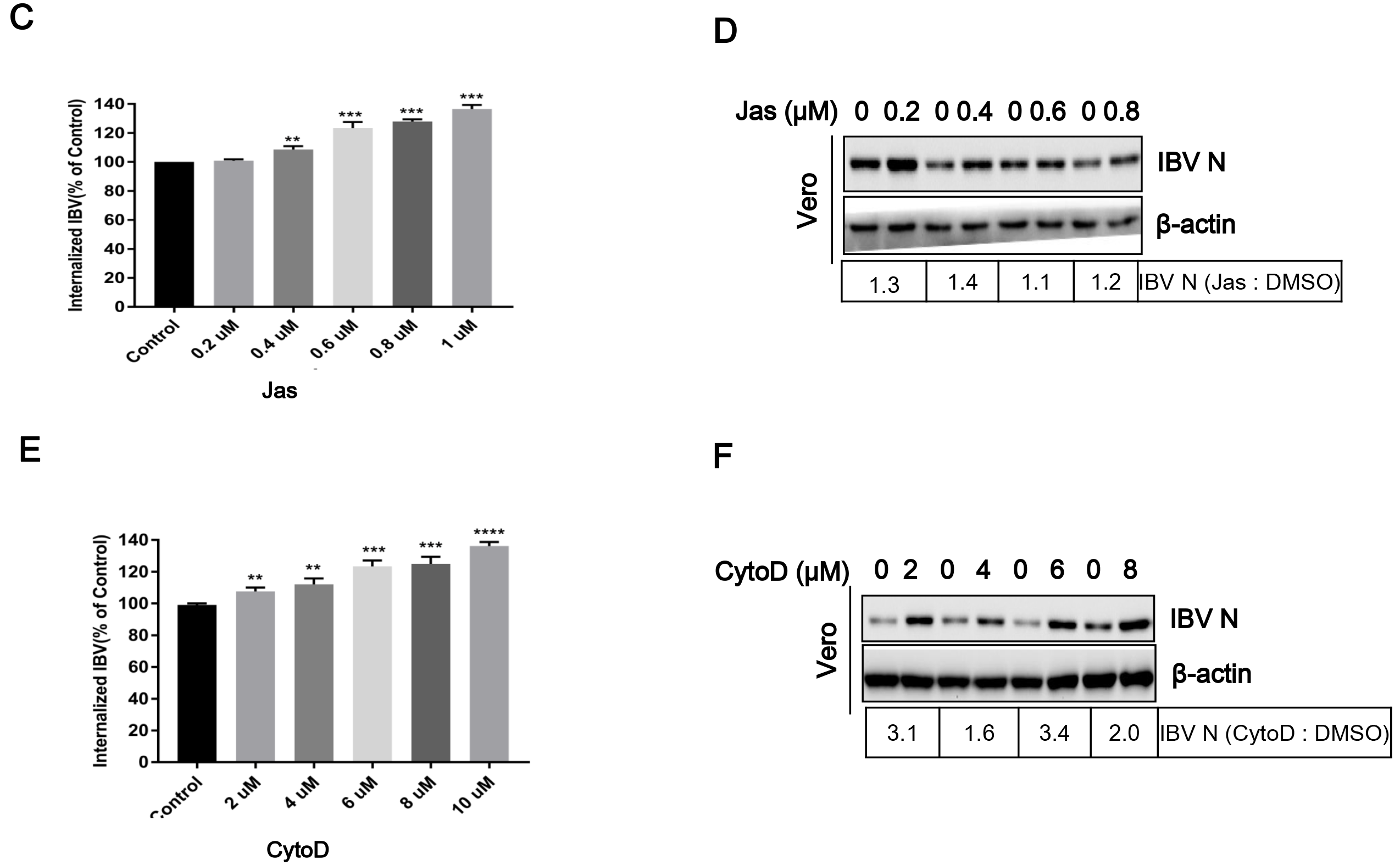
IBV entry causes actin rearrangement. (A) Vero cells were incubated with IBV (MOI 5) at 4°C for 1 h to allow virus attachment, and the unbound virus particles were washed away by DMEM. Mock infection was set as control group. Temperature was shifted to 37°C to allow virus entry. Cells were fixed at 15, 30, 45, and 60 min.p.i., and incubated with Alexa Fluor 488-phalloidin (green) for 1 h to stain actin filaments, followed with IBV N protein staining (red) and DAPI staining (blue). The signals were observed under LSM880 confocal laser-scanning microscope (Zeiss). Representative images were shown. Scale bars=20 μm. (B) Vero cells were infected with R18-IBV (MOI 5) at 4°C for 1 h, and temperature was shifted to 37 °C to allow internalization. R18-mock infection was set as control group. Cells were fixed at indicated time points and incubated with Alexa Fluor 488-phalloidin to stain actin filaments (green). Cell nuclei were stained with DAPI (blue). The signals were observed under LSM880 confocal laser-scanning microscope (Zeiss). Representative images were shown. Scale bars=20 μm. Red signal: R18-IBV; green signal: actin; blue signal: nuclei. (C-F) Vero cells were pretreated with increasing concentrations of CytoD or Jas for 30 min and infected with IBV (MOI 5). The internalized viral gRNA was assayed at 2 h.p.i. (C and E), and the expression of IBV N protein was determined at 8 h.p.i. (D and F). The experiment was performed in triplicate, and average values with stand errors were presented in panel C and E. The intensity of IBV N was determined with Image J, normalized to β-actin, and shown as fold change of Jas: DMSO or CytoD: DMSO in table panel of D and F.

We further investigated the role of the actin cytoskeleton in IBV entry by using chemical inhibitors to interfere with actin polymerization or de-polymerization. Vero cells were treated with jasplakinolide (Jas), an actin polymer-stabilizing drug (74–76), or with cytochalasm D (CytoD), which binds to the growing ends of actin filaments and prevents the polymerization (77). Vero cells were pretreated with either Jas or CytoD in increasing concentrations before being infected with IBV. Cells were harvested at indicated time points to measure IBV gRNA internalization or N protein expression. Fig. 8C-D showed that when actin polymers were stabilized by Jas, the IBV internalization and N protein expression were increased in a dose-dependent manner. Fig. 8E-F showed that when CytoD prevented the formation of actin polymers, IBV internalization and N protein expression were also significantly increased. This observation is self-contradictory and out of expectation. Thus, we use the well characterized VSV as control to perform this experiment, ensuring the inhibitors work. When the Jas or CytoD was applied to Vero cells before VSV infection, both inhibitors reduced VSV G protein expression (Supplementary Fig. 2B, C), consistent with previous report of actin dependent entry of VSV (73). These results imply that the inhibitors indeed interfere with actin polymerization or de-polymerization. The enhancement of IBV infection by these inhibitors is not the only case, it has been reported that SFV infection was also enhanced by CytoD pretreatment (78). In all, above results demonstrate that the binding of IBV to cell surface causes actin de-polymerization and rearrangement, however, when the actin de-polymerization and rearrangement are interfered by inhibitors, the virus entry is enhanced. How do the inhibitors increase IBV entry? The underlying mechanisms are unclear.

### IBV moves across the entire endo-lysosomal system

After identification of the endocytic pathway hijacked by IBV, we next tracked the transport vesicles utilized by incoming IBV. R18-IBV was incubated with Vero cells at 4°C for 1 h, allowing attachment. R18-mock infected cells were included as control. After removing the unbound virus by replacing with fresh DMEM, the temperature was shifted to 37°C to facilitate the synchronous endocytosis. Cells were subjected to immunostaining by using antibodies against early endosome marker Rab5, late endosome marker Rab7, and lysosome marker LAMP1. Rab5 and Rab7 are GTPases that are required for the movement of endocytosed cargo to early or late endosomes (79, 80). The co-localization of R18-IBV and transport vesicles markers at 0-3 h.p.i. was observed under confocal microscope. As shown in Fig. 9A, there was no red signal detected in R18-mock infected cells ( first panel); the overlapped signals of R18-IBV and early endosome marker Rab5 were mainly observed at 1 h.p.i. (yellow dots with white arrows in second panel); however, there was no co-image of R18-IBV with Rab5 at 2 h.p.i. and 3 h.p.i.(third and fourth panels). This reveals that IBV enters into early endosome at 1 h.p.i, and leaves early endosome at 2 and 3 h.p.i.. Moreover, the co-localization of virus particles and Rab7 was mainly observed at 2 h.p.i. (yellow dots with white arrow in third panel), there was no R18-IBV overlapped with late endosome marker Rab7 observed at 1 h.p.i. and 3 h.p.i., demonstrating that R18-IBV enters late endosome at 2 h.p.i.. Furthermore, R18-IBV was found in LAMP1 containing lysosomes at 3 h.p.i. (yellow dots with white arrow in fourth panel). Previous study showed that VSV was internalized into early endosome in a few minutes via CME (81). Thus, we used R18-VSV to perform above experiment in Vero cells as control. As shown in Supplementary Fig. 3, R18-VSV co-localized well with early endosome marker Rab5 at 5, 10, and 20 min.p.i. (yellow dots with white arrows in second, third, and fourth panels). There was less co-image of R18-VSV with Rab5 at 30 min.p.i. (fifth panel), indicating the R18-VSV leaves early endosomes at this time point. Moreover, there was almost no co-image of R18-VSV with Rab7/LAMP1 at 0-30 min. These observations verify that VSV enters early endosome, but is not delivered into late endosome and lysosome, consistent with previous study (73). In all, above results reveal that IBV enters early endosome at around 1 h.p.i., passes through late endosome at around 2 h.p.i., and reaches lysosome at 3 h.p.i..

**Figure 9.**
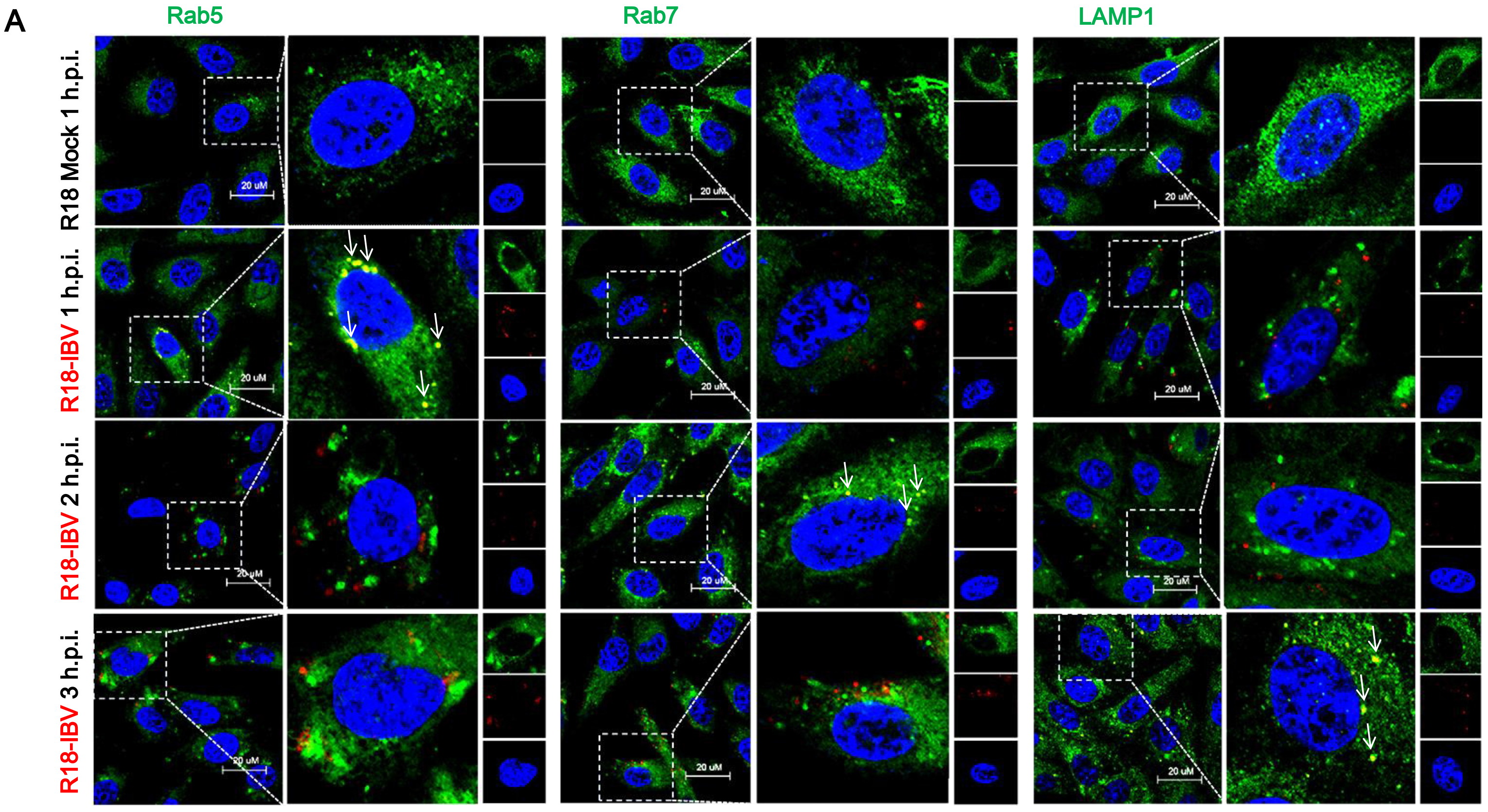
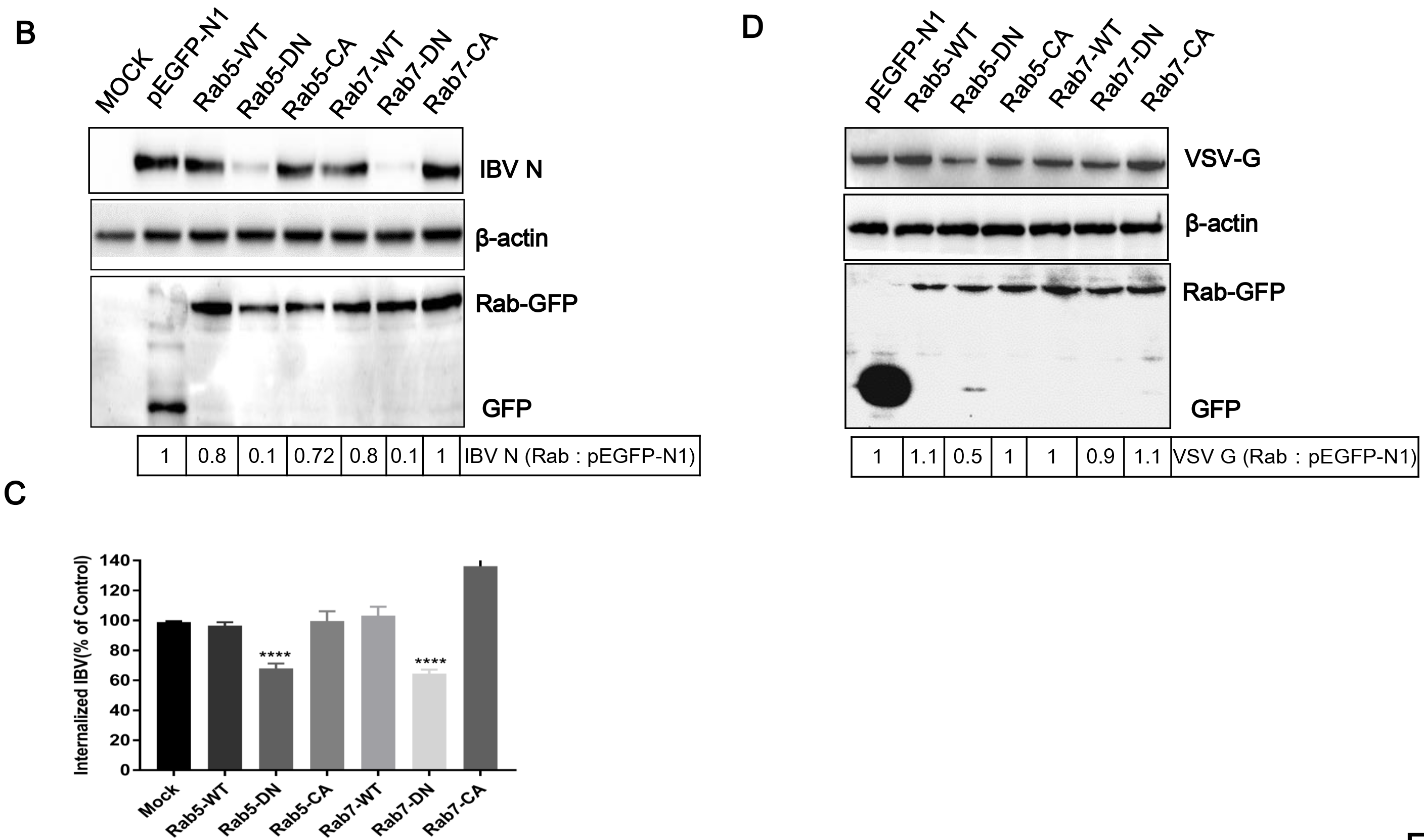
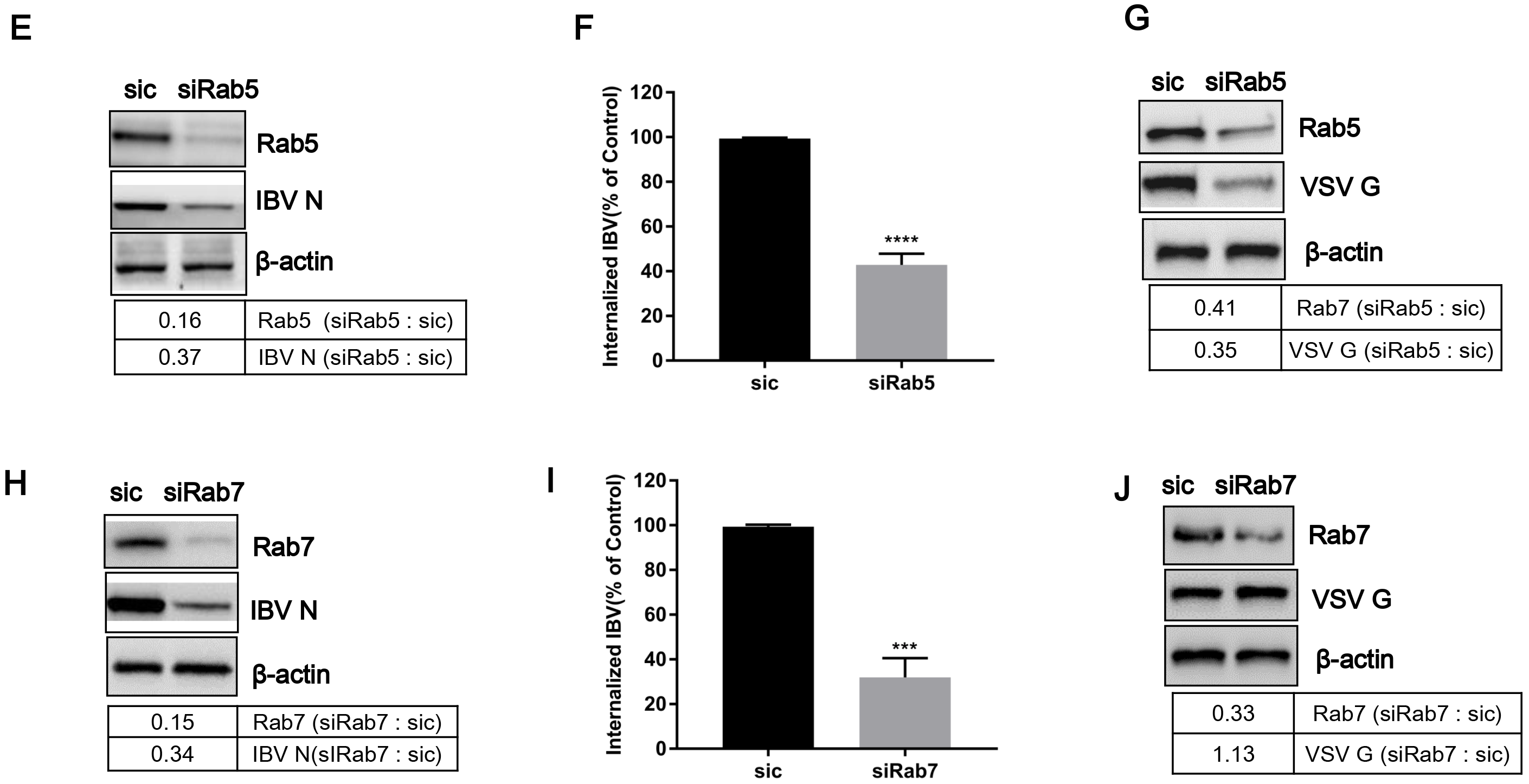
The incoming R18-IBV transports along the entire endocytic pathway. (A) R18-IBV infected Vero cells (MOI 5) were immunostained with Rab5, Rab7, or LAMP1 at 1, 2, 3 h.p.i., respectively, followed with DAPI staining. R18-mock infection was included in a parallel experiment as control. The signals were observed under LSM880 confocal laser-scanning microscope (Zeiss). Representative images were shown. Scale bars=20 μm. Red signal: R18-IBV; green signal: early endosome (Rab5), later endosome (Rab7), or lysosome (LAMP1); blue signal: nuclei. (B-D) Cells were transiently transfected with constructs encoding GFP, GFP-Rab5-WT, GFP-Rab5-DN, GFP-Rab5-CA, GFP-Rab7-WT, GFP-Rab7-DN, and GFP-Rab7-CA for 24 h, followed with IBV infection (H1299 cells) or VSV infection (Vero cells). The expression of IBV N protein, VSV G protein, GFP, GFP-Rab5, and GFP-Rab7 were detected at 8 h.p.i. (B, D); the incoming virus level was determined by quantification of viral gRNA at 2 h.p.i. (C). The experiment was performed in triplicate, and average values with stand errors were presented in panel C. (E-J) Cells were transfected with siRNAs targeted to either Rab5 or Rab7, respectively. Non-target siRNA (sic) was transfected as control. At 36 h post-transfection, cells were infected with IBV (H1299 cells) or VSV (Vero cells). The knock down level of Rab5/Rab7 and the expression of IBV N or VSV G was detected by Western Blot analysis at 8 h.p.i. (E, G; H, and J). The incoming IBV level was determined by quantification of viral gRNA at 2 h.p.i. (F and I). The experiment was performed in triplicate, and average values with stand errors were presented in panel F and I. The intensity of IBV N or VSV G was determined with Image J, normalized to β-actin, and shown as fold change of Rab: pEGFP-N1, siRab5: sic, or siRab7: sic in table panel of B, D, E, G, H, and J..

As IBV co-localized extensively with endo-lysosomal compartments, we asked whether all these vehicles are really involved in IBV infectious route. To achieve this, we took advantage of the fact that, in addition to providing organelle identity, Rab5 and Rab7 serve critical roles in regulation of cargo transport within the endosomal system (82). Accordingly, dominant negative mutant of these proteins has been shown to block cargo transport (83, 84). Expression of Rab5-DN prevents fusion of endocytic vesicles with early endosomes, while Rab7-DN prevents movement from early to late endosomes. H1299 cells were transfected with GFP, GFP fused Rabs-WT (Rab5-WT, Rab7-WT), Rabs-DN (GFP-Rab5-S34N, GFP-Rab7-T22N), or Rabs-CA (GFP-Rab5-Q79L, GFP-Rab7-Q67L), and then the cells were infected with IBV for 12 h. The cells were harvested for Western blot to check the expression of Rabs and IBV N protein. As shown in Fig. 9B, all these proteins were successfully expressed. Compared with GFP expressing group, expression of Rab5-WT, Rab5-CA, Rab7-WT, and Rab7-CA has only minimal effect on the expression of IBV N, whereas expression of Rab5-DN and Rab7-DN caused significantly reduction (0.1-fold) in IBV infection (Fig. 9B). These data reveal that IBV could not infect cells expressing Rab5-DN and Rab7-DN. Inhibition of IBV infection by Rab5-DN and Rab7-DN was highly significant, indicating that transport through the early and late endosome is necessary for IBV infection. To further verify above observation, we quantified the internalized IBV in Rabs expressing cells. Results in Fig. 9C confirmed that expression of Rab5-DN and Rab7-DN decreased internalization of IBV gRNA by 0.6-fold. Moreover, expression of Rab7-CA slightly increased virus internalization. To ensure our Rabs construct was functional, we examine the effect of Rabs on VSV infection in Vero cells. As shown previously (73), Rab5-DN, but not Rab7-DN, inhibited VSV infection (Fig. 9D). Collectively, these data indicate that major IBV virions are delivered to early endosomes after CME, followed by transferring to late endosomes.

Knock down of Rab5 or Rab7 was carried out to confirm above conclusion. H1299 cells were transfected with siRab5 or siRab7, followed with IBV infection. The knock down effect of Rabs and IBV N protein expression was examined by Western blot. Fig. 9E and 9H showed that both Rab5 and Rab7 expression were successfully knocked down by RNA interference in H1299 cells, resulting lower expression of IBV N protein than that in sic tranfected cells. This demonstrates that the virus replication is decreased when the transport pathway is hampered by removing Rab5 or Rab7. Quantification of incoming virus genome also confirmed that depletion of Rab5 or Rab7 reduced virus genome internalization (Fig. 9 F and I). To ensure siRabs was functional, we examine the effect of Rabs knock down on VSV infection in Vero cells. Western blot result showed that knock down of Rab5 reduced VSV G protein expression (Fig. 9G), while knock down of Rab7 did not inhibit VSV infection (Fig. 9J). Altogether, these data further confirm that after internalized by CME, both early and late endosomes are required for IBV transport.

### IBV fuses with late endosomes or lysosomes

To understand the fusion of IBV envelope with cellular membranes, we labeled IBV virion with two fluorescent lipids, R18 (red) and DiOC (green), a labeling method developed by Sakai and coworkers (85). The intact virus membrane displays the red color as the high concentration of R18 quenches the fluorescence signal emitted by the DiOC. When the virus membrane fuses with cellular membrane, the two lipids were diluted and the green fluorescence of DiOC was no longer quenched by R18, red and green color will be displayed respectively. The Vero cell culture supernatant was labeled with R18 and DiOC as mock infection sample. Vero cells were infected with R18/DiOC-IBV or with R18/DiOC-mock for 1, 2, 3 h, stained with Rab5, Rab7, LAMP1 antibodies, and observed under confocal microscope. As shown in Fig. 10, in mock infection group, there was neither red signal nor green signal observed (first panel). When the R18/DiOC labeled virus was incubated with cells, only red color spots were observed at 1 and 2 h.p.i., no green signals were detected at these two time points (second and third panels). Moreover, most of the red spots were co-localized with early endosomes marker Rab5 at 1 h.p.i. (pink spots with white arrows in second panel), and transferred to Rab7 containing late endosomes or LAMP1 containing lysosomes at 2 h.p.i. (pink spots with white arrows in third panel). Gradually, green spots (membrane fusion signals) were observed and increased in numbers at 3 h.p.i. (white arrow in fourth panel), overlapped with either late endosome marker Rab7 or lysosome marker LAMP1 (white arrows in fourth panel), implying that IBV membranes undergo fusion with late endosome/lysosome membranes at 3 h.p.i. or between 2-3 h.p.i.. We also performed above fusion experiment by using well studied VSV labeled with R18/DiOC as control, which has been reported to fuse with early endosomes. As shown in Supplementary Fig.4, only red spots were observed at 10 min.p.i., overlapped with Rab5 (yellow spots with white arrows in second panel), without overlap with Rab7 or LAMP1. At 20 and 30 min.p.i., green spots were detected, co-localized with Rab5 (pink spots with white arrows in third and fourth panels), consistent with previous report (73). In all, we demonstrate that IBV fuses with late endosome/lysosome at 2-3 h.p.i., and its intracellular trafficking is slower than VSV.

**Figure 10.**
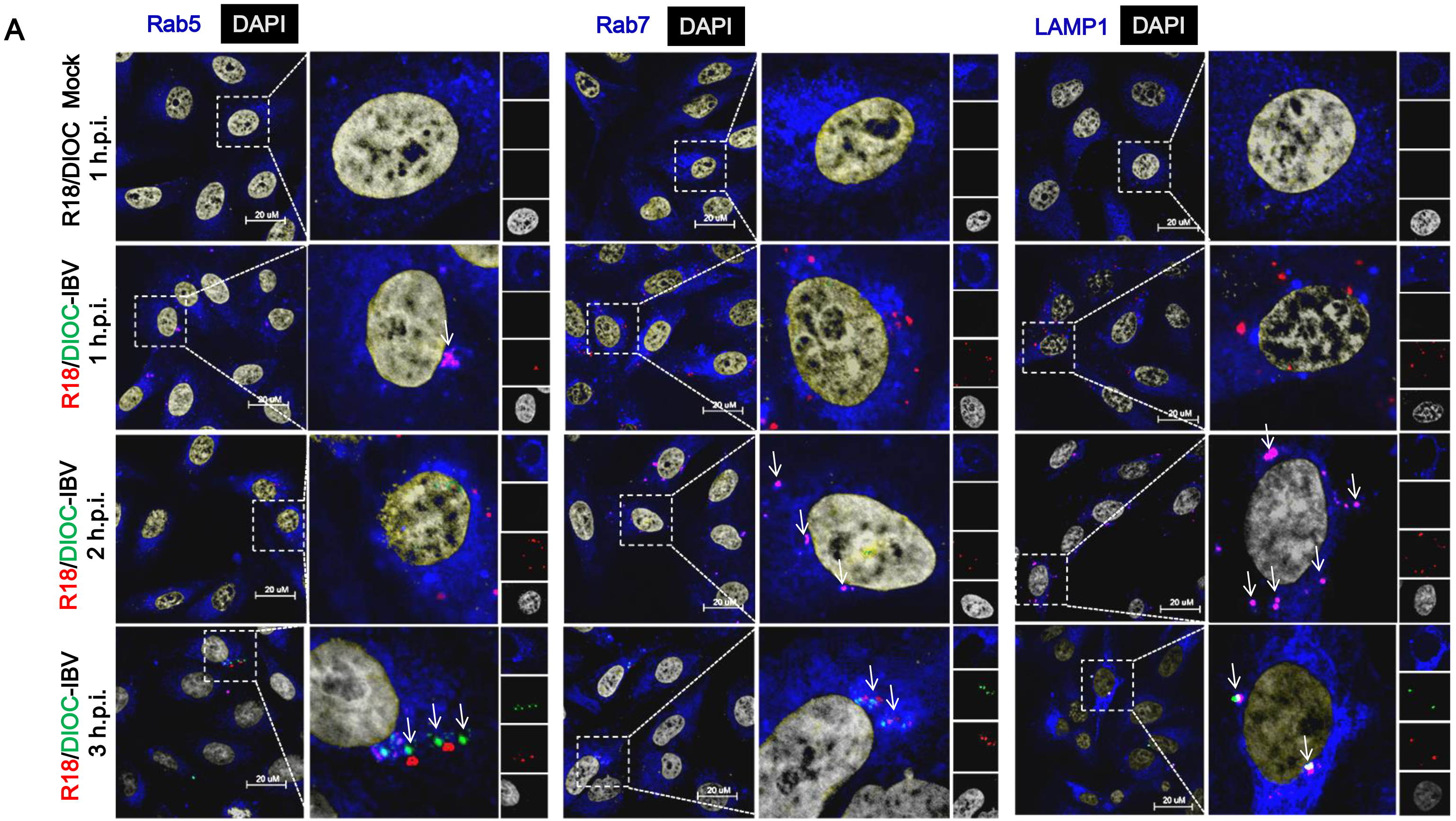
IBV fuses with late endosomes or lysosomes membranes. Vero cells were infected with R18-DiOC-IBV (MOI 5) for 1, 2, 3 h, and subjected to immunostaining using Rab5, Rab7, or LAMP1 antibody. The separation of R18 (red) and DiOC (green) signal, and co-localization with vehicle markers (blue) were observed under LSM880 confocal laser-scanning microscope (Zeiss). Representative images were shown. Scale bars=20 μm. Red signal: R18-IBV; green signal: DiOC-IBV; Blue Signal: early endosome (Rab5), later endosome (Rab7), or lysosome (LAMP1); Grey signal: nuclei.

### Valosin-Containing Protein (VCP) is involved in IBV trafficking

Our results so far demonstrate that IBV pass through early and late endosomes. Thus, maturation of endosome should provide the environmental cues required for productive virus-cell fusion. VCP is an abundant protein implicated in an increasing number of biological processes, including membrane fusion after mitosis (86), retro-translocation of unfolded proteins from the endoplasmic reticulum (87, 88), spindle assembly (89), and endosome trafficking (90, 91). Previous report shows that depletion of VCP results in an accumulation of virus particles in the early endosomal compartments, suggesting that VCP is required for the maturation of endosomes and efficient transfer of virus particles from the early to late endosomes (92). To further access the role of VCP in IBV infection, we constructed GFP-tagged VCP and transiently expressed this protein in H1299 cells, followed with infection. The expression of GFP-VCP was confirmed by Western blot (Fig. 11A). Compared with the GFP expressing control group, overexpression of GFP-VCP significantly increased the expression of IBV N by 1.45-fold (Fig. 11A), however, the virus internalization was not increased (Fig. 11B), indicating VCP plays a role post-internalization. The higher rate of viral gRNA replication associated with VCP expression was also detected with RT-PCR by using primers targeting to negative stranded gRNA (Fig. 11C). Immunofluorescence in Fig.11D showed that GFP-VCP was co-localized with Rab5 (yellow in second panel), instead of Rab7 or LAMP1, suggesting the role of VCP in early endosome maturation. Taken together, these results demonstrate that VCP is involved in transferring IBV from early endosomes to destination for efficient replication.

**Figure 11.**
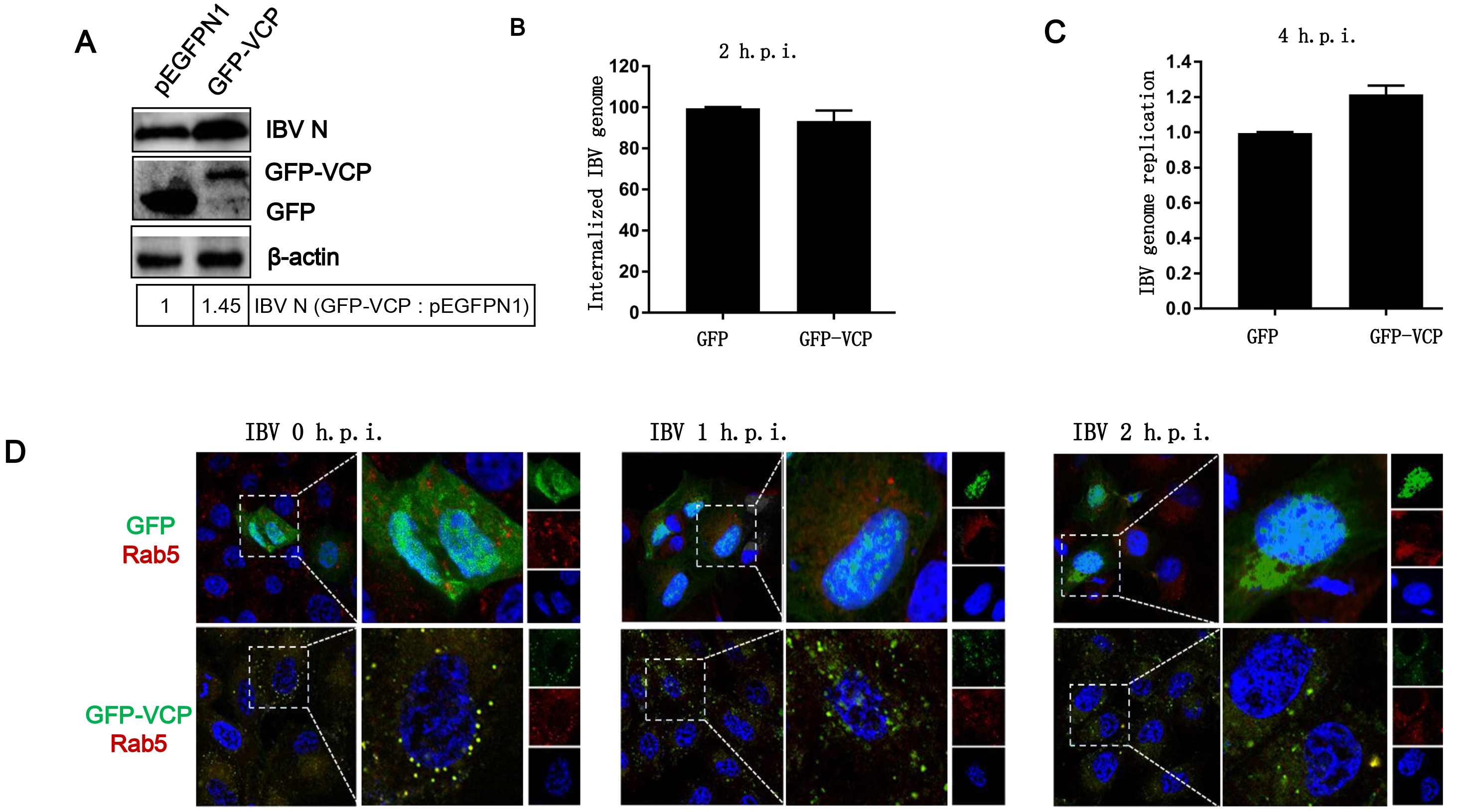
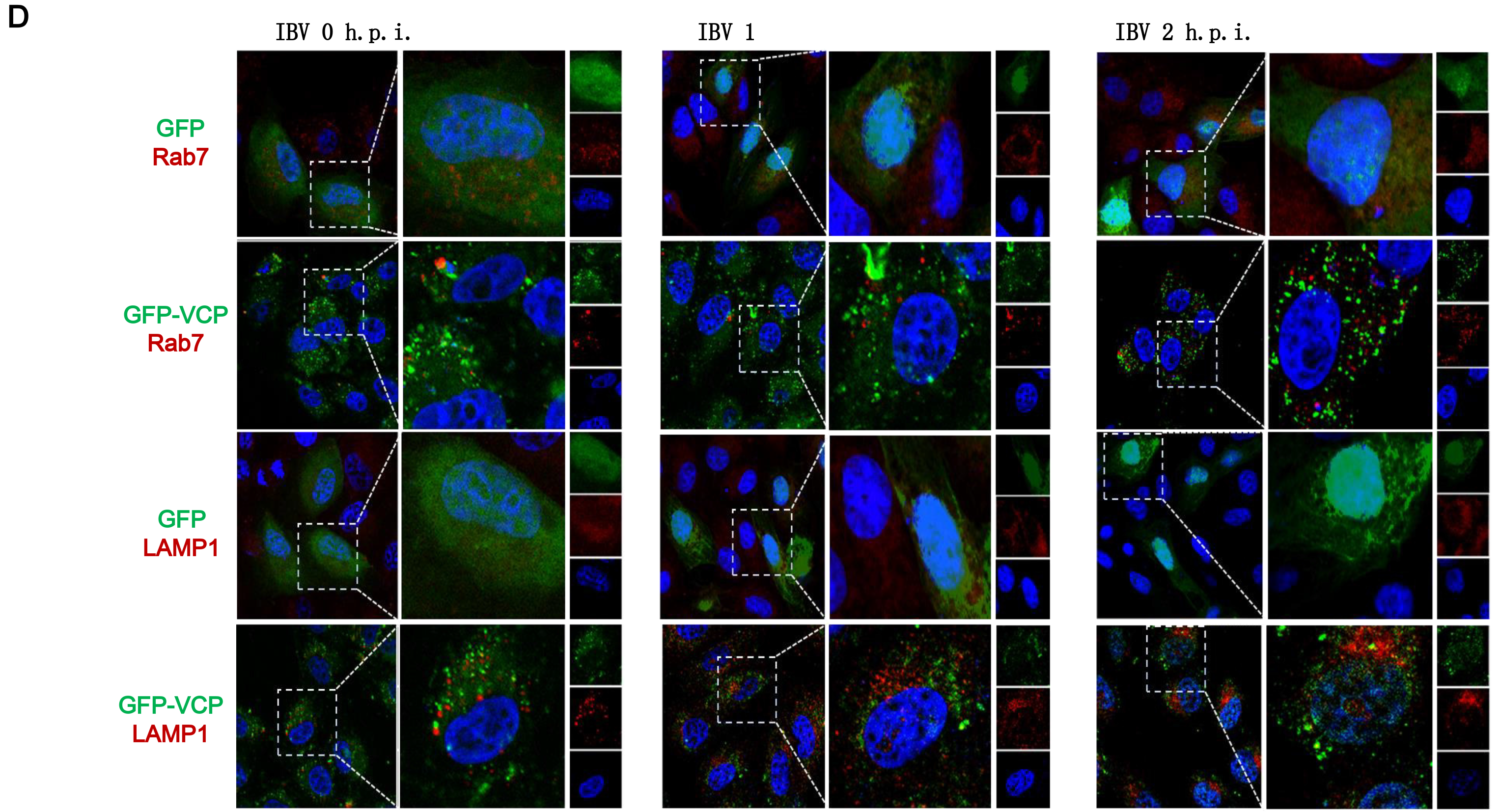
VCP is involved in IBV trafficking. H1299 cells were transfected with constructs expressing GFP-VCP or vector pEGFP-N1 for 24 h, followed by IBV Infection. (A) The expression level of GFP, GFP-VCP and IBV-N was scored by Western blot at 8 h.p.i.. The band intensity of IBV N was determined with Image J, normalized to β-actin, and shown as fold change of GFP-VCP: pEGFPN1 in table panel.. (B) The internalized IBV gRNA was analyzed with semi-quantitative real time RT-PCR at 2 h.p.i.. (C) The replication level of IBV gRNA was determined by semi-quantitative real time RT-PCR with primers targeting to negative stranded gRNA at 4 h.p.i. The experiment was performed in triplicate, and average values with stand errors were presented in panel B and C. (D) Co-image of GFP-VCP with Vesicle markers (Rab5, Rab7, and LAMP1) was determined by immunofluoresence at 0, 1, and 2 h.p.i.. GFP was set as control. Representative images were shown. Scale bars=20 μm. Green signal: GFP or GFP-VCP; Red signal: Rab5, Rab7, or LAMP1.

## DISCUSSION

Viruses must enter the host cell to start their life cycle. Thus, characterization of viral entry pathway is important for understanding of virus pathogenesis and design of anti-viral drugs. Envelope viruses penetrate cells by two different ways: fusion with cell plasma membrane or receptor-mediated endocytosis. Direct membrane fusion on the cell surface is independent of low pH, and access through the endocytosis pathway usually relies on the low pH of the endocytic vesicles. Different endocytosis pathways have been characterized over the past decade, in which the CME is one of the most commonly used by many envelope viruses (93). Coronavirus entry is mediated by S protein, which is responsible for receptor binding and membrane fusion. Previously, it has been demonstrated that SARS-CoV is internalized by CME in human hepatoma HepG2 cell line (39), however, in HEK293E-ACE2-GFP cells, the entry of SARS-CoV is clathrin- and caveolae-independent (40); MHV entry is mediated by CME through the endo-/lysosomal pathway in proteolysis-dependent manner (43, 94-97); PHEV enters into cells via CME in Rab5, cholesterol, and low pH dependent manner (98); HCoV-NL63 requires CME for successful entry into the LLC-MK2 cells (99); for the HCoV-229E, a CavME uptake has been suggested (16), however, the clinical isolates of HCoV-229E bypass the endosome for cell entry (100). Above evidence demonstrates that coronavirus enter cells via multiple pathways. Thus, study on IBV entry mechanisms is necessary for understanding group 3 coronavirus infection process. As mentioned previously, the infectivity of IBV on BHK cells is inhibited by NH_4_Cl treatment (44), demonstrating that IBV entry requires low pH triggered membrane fusion. In this study, we made an effort to systematically examine every step of this process. First, we found that IBV particles initially attach to lipid rafts on plasma membrane, and enter cells via CME, which requires the help of CHC and dynamin1. Additionally, we demonstrate that IBV particles are transported via early and late endosome, and fuse with late endosome-lysosome membrane to release the nucleocapsid.

Lipid rafts are involved in regulation of different biological events, serve as organizing centers of signaling molecules assembly, and influence membrane fluidity and membrane protein trafficking (101). The involvement of lipid rafts in virus entry, assembly and budding has been demonstrated by either the co-localization study of the viral structural proteins and lipid rafts or the effects of lipid rafts disrupting agents on the replication processes of several viruses (102). The best characterized lipid rafts-related non-enveloped viral entry is simian virus 40 (SV40) and echovirus type 1 (EV1) (103, 104). Binding of SV40 with MHC class I receptors triggers receptor clustering and redistribution, more caveolae is recruited from the cytoplasm at the site of entry, resulting in caveolae-mediated endocytosis in about 20 min (104). In some cell types, SV40 can enter the caveosomes directly from lipid rafts in non-coated vesicles (105). Similar to SV40, EV1 attachment and binding with cells triggers clustering and relocation of α2β1-integrin receptors from lipid rafts to the caveolae-like structures (104). Depletion of cholesterol in lipid rafts inhibits EV1 infection (103). For envelope virus, Semliki Forest virus (SFV) and Sindbis virus (SIN) require cholesterol and sphingolipids in target membrane lipid rafts for envelope glycoprotein-mediated membrane fusion and entry (29). Many envelope virus receptors are located in lipid rafts or would be relocated into lipid rafts after infection, such as Human T-lymphotropic virus Type I (HTLV-1) receptor glucose transporter 1 (GLUT-1), Ebola virus and Marburg virus folate receptor-α (FRα), Hepatitis B virus complement receptor type 2 (CR2), Human herpesvirus 6 (HHV-6) receptor CD46 (106). An alternative receptor for Human Immunodeficiency virus (HIV-1) envelope glycoprotein on epithelial cells is glycosphingolipid galactosyl-ceramide (GalCer), which also enriches at lipid rafts (107, 108). Membrane cholesterol is also involved in the first stages of flavivirus infection (47, 109). Guo *et al* demonstrate that IBV structural proteins, but not non-structural proteins, migrate and integrate into lipid rafts after viral protein translation. The cholesterol enriched microdomains were only involved in IBV attachment, not virus replication or virus particle release (38). However, the involvement mechanisms of lipid rafts on IBV infectivity are still unclear. In our study, we substantiated the involvement of lipid rafts on virus life cycle via disruption of lipid rafts by using chemicals at pre-, during-, or post-infection, and checked IBV internalization, viral protein expression, and progeny virus release. Results showed that lipid rafts were involved in IBV entry step. Cholesterol replenishment experiment further confirmed that lipid rafts were necessary for IBV entry. Membrane flotation assay showed the co-fractionation of IBV particles and lipid rafts during absorption step. Immunofluoresence showed that R18-IBV associates with lipid rafts on the cells surface. Above evidence substantiates that IBV initiates the entry step by attaching to lipid rafts.

Further experiment showed that chemical inhibitor of endosome acidification (NH_4_Cl) treatment in various susceptible cells reduced the IBV internalization and protein expression, suggesting a requirement for transport of virions to endosomes. To further confirm our observations, we used inhibitor CPZ to interfere with CME, or knock down of CHC siRNA to specifically block CME, and we found that IBV internalization and gene expression were significantly decreased. This evidence determines the importance of CME in IBV entry, which is widely hijacked by various envelope viruses. Moreover, the importance of dynamin1, a GTPase that helps to snap the vesicle from plasma membrane, was determined during IBV entry. Lipid rafts is often considered to be involved in CavME. To explain these contradictory observations, Huh7, a caveolin deficient cell line (110, 111), was used in the cholesterol depletion experiment. When cholesterol was depleted by chemicals or replenished in Huh7 cells, similar inhibition effect on IBV entry was observed as that in Vero and H1299 cells. These results confirm that although IBV attachment and entry depend on the intact lipid rafts, CME is the main endocytotic pathway for internalization.

Our final research question was the virus trafficking route. By using R18-labeled IBV to infect cells, we found that IBV attachment leads to actin rearrangement and co-localized with actin filament after internalization. This indicates that virus internalization requires the actin de-polymerization or polymerization, to facilitate the virus containing vesicles to move along actin filament. However, stabilization of the actin filament using Jas and inhibition of actin polymerization using CytoD resulted in increase of virus entry. Rab5 and Rab7 GTPases are the key regulators of transport to early and late endosomes, and ablation of Rab5 and Rab7 has been widely used to study viral entry. Using R18-labeled virus, we found that after internalization, incoming IBV particles were first co-localized with Rab5 and then with Rab7, finally overlapped with lysosome marker LAMP1, across the entire endocytic network. The importance of Rab proteins were examined by overexpression and RNA interference technology. The results showed that Rab5 and Rab7 played crucial roles for IBV entry, confirming that IBV trafficking really depends on early and late endosomes. Also, using the R18 and DiOC double labeled virus, we revealed that the membrane fusion happens at 2-3 h.p.i., the time point of virus reaches late endosome-lysosome. Overexpression of VCP, a protein involved in endosome maturation, increased virus gene expression. We speculated that IBV particles were first transported to early endosomes in Rab5-demepdent manner, and were guided to late endosomes in Rab7 and VCP dependent manner. Also, the Rab7 dependence indicates that membrane fusion depends on endosomal maturation, suggesting that low pH environment in the late endosome-lysosome may provide an important signal for fusion and viral gRNA release.

In conclusion, we conduct a systematic study to dissect IBV entry process in susceptible cell lines. The evidence presented here indicates that IBV attaches to lipid rafts, induces actin de-polymerization, internalizes into clathrin coated vesicles via dynamin1 snapping, transports along early/late endosome, and lysosome, finally fuses with late endosome-lysosome membrane, releases genome in to cytoplasm (Fig. 12). An understanding of the cellular components involved in IBV invasion into the cell, is likely to open new opportunities to develop novel therapeutic approaches.

**Figure 12.**
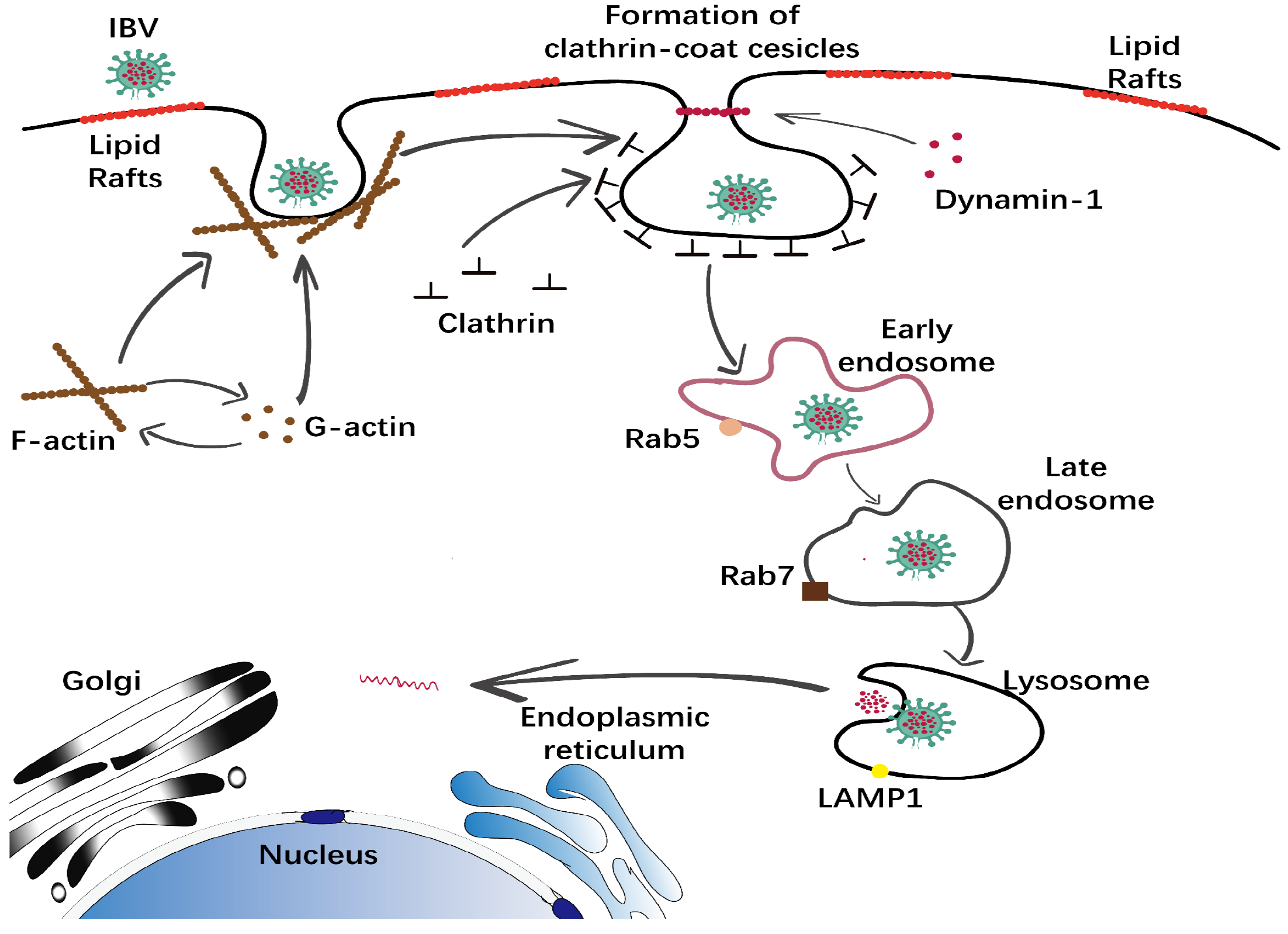
Model of IBV internalization, trafficking, and fusion. IBV attaches to lipid rafts on the cell surface and is internalized from the plasma membrane via CME, with the help of dynamin1 and actin de-polymerization. The virus containing clathrin coated vesicles were taken up in Rab5-containing early endosomes, transferred into late endosomes and lysosomes. The low pH environment in late endosomes-lysosomes induces virus-cell membrane fusion.

## ACKNOWLEDGEMENTS

This study was supported by National Natural Science foundation of China (Grant No. 31772724), Chinese Special Fund for Agricultural Sciences Research in the Public Interest (Grant No. 2014JB16; Grant No. 2017JB05), China Ministry of Science and Technology (Grant No. 2017YFD0500802), Elite Youth Program of Chinese Academy of Agricultural Sciences (Grant No. 20170260401), and National Natural Science foundation of China (Grant No. 31530074).

We acknowledge the gifts of IBV Beaudette strain and viral protein antibodies from Prof. Liu Dingxiang’s lab in South China Agricultural University, China.

## REFERENCES

1. Cossart P, Helenius A. 2014. Endocytosis of viruses and bacteria. Cold Spring Harb Perspect Biol 6.

2. Yamauchi Y, Helenius A. 2013. Virus entry at a glance. J Cell Sci 126:1289–95.

3. Suomalainen M, Greber UF. 2013. Uncoating of non-enveloped viruses. Curr Opin Virol 3:27–33.

4. Harrison SC. 2015. Viral membrane fusion. Virology 479–480:498–507.

5. Keen JH. 1990. Clathrin and associated assembly and disassembly proteins. Annu Rev Biochem 59:415–38.

6. Kirchhausen T. 1999. Adaptors for clathrin-mediated traffic. Annu Rev Cell Dev Biol 15:705–32.

7. Cureton DK, Massol RH, Whelan SP, Kirchhausen T. 2010. The length of vesicular stomatitis virus particles dictates a need for actin assembly during clathrin-dependent endocytosis. PLoS Pathog 6:e1001127.

8. Meier O, Boucke K, Hammer SV, Keller S, Stidwill RP, Hemmi S, Greber UF. 2002. Adenovirus triggers macropinocytosis and endosomal leakage together with its clathrin-mediated uptake. J Cell Biol 158:1119–31.

9. Hommelgaard AM, Roepstorff K, Vilhardt F, Torgersen ML, Sandvig K, van Deurs B. 2005. Caveolae: stable membrane domains with a potential for internalization. Traffic 6:720–4.

10. Insel PA, Head BP, Ostrom RS, Patel HH, Swaney JS, Tang CM, Roth DM. 2005. Caveolae and lipid rafts: G protein-coupled receptor signaling microdomains in cardiac myocytes. Ann N Y Acad Sci 1047:166–72.

11. Pelkmans L, Kartenbeck J, Helenius A. 2001. Caveolar endocytosis of simian virus 40 reveals a new two-step vesicular-transport pathway to the ER. Nat Cell Biol 3:473–83.

12. Marjomaki V, Pietiainen V, Matilainen H, Upla P, Ivaska J, Nissinen L, Reunanen H, Huttunen P, Hyypia T, Heino J. 2002. Internalization of echovirus 1 in caveolae. J Virol 76:1856–65.

13. Bousarghin L, Touze A, Sizaret PY, Coursaget P. 2003. Human papillomavirus types 16, 31, and 58 use different endocytosis pathways to enter cells. J Virol 77:3846–50.

14. Empig CJ, Goldsmith MA. 2002. Association of the caveola vesicular system with cellular entry by filoviruses. J Virol 76:5266–70.

15. Beer C, Andersen DS, Rojek A, Pedersen L. 2005. Caveola-dependent endocytic entry of amphotropic murine leukemia virus. J Virol 79:10776–87.

16. Nomura R, Kiyota A, Suzaki E, Kataoka K, Ohe Y, Miyamoto K, Senda T, Fujimoto T. 2004. Human coronavirus 229E binds to CD13 in rafts and enters the cell through caveolae. J Virol 78:8701–8.

17. Kalin S, Amstutz B, Gastaldelli M, Wolfrum N, Boucke K, Havenga M, DiGennaro F, Liska N, Hemmi S, Greber UF. 2010. Macropinocytotic uptake and infection of human epithelial cells with species B2 adenovirus type 35. J Virol 84:5336–50.

18. Inal JM, Jorfi S. 2013. Coxsackievirus B transmission and possible new roles for extracellular vesicles. Biochem Soc Trans 41:299–302.

19. de Vries E, Tscherne DM, Wienholts MJ, Cobos-Jimenez V, Scholte F, Garcia-Sastre A, Rottier PJ, de Haan CA. 2011. Dissection of the influenza A virus endocytic routes reveals macropinocytosis as an alternative entry pathway. PLoS Pathog 7:e1001329.

20. Aleksandrowicz P, Marzi A, Biedenkopf N, Beimforde N, Becker S, Hoenen T, Feldmann H, Schnittler HJ. 2011. Ebola virus enters host cells by macropinocytosis and clathrin-mediated endocytosis. J Infect Dis 204 Suppl 3:S957–67.

21. Rizopoulos Z, Balistreri G, Kilcher S, Martin CK, Syedbasha M, Helenius A, Mercer J. 2015. Vaccinia Virus Infection Requires Maturation of Macropinosomes. Traffic 16:814–31.

22. Vogt C, Eickmann M, Diederich S, Moll M, Maisner A. 2005. Endocytosis of the Nipah virus glycoproteins. J Virol 79:3865–72.

23. Raghu H, Sharma-Walia N, Veettil MV, Sadagopan S, Chandran B. 2009. Kaposi’s sarcoma-associated herpesvirus utilizes an actin polymerization-dependent macropinocytic pathway to enter human dermal microvascular endothelial and human umbilical vein endothelial cells. J Virol 83:4895–911.

24. Tan L, Zhang Y, Zhan Y, Yuan Y, Sun Y, Qiu X, Meng C, Song C, Liao Y, Ding C. 2016. Newcastle disease virus employs macropinocytosis and Rab5a-dependent intracellular trafficking to infect DF-1 cells. Oncotarget 7:86117–86133.

25. Simons K, Ikonen E. 1997. Functional rafts in cell membranes. Nature 387:569–72.

26. Fivaz M, Abrami L, van der Goot FG. 1999. Landing on lipid rafts. Trends Cell Biol 9:212–3.

27. Simons K, Toomre D. 2000. Lipid rafts and signal transduction. Nat Rev Mol Cell Biol 1:31–9.

28. Kovbasnjuk O, Edidin M, Donowitz M. 2001. Role of lipid rafts in Shiga toxin 1 interaction with the apical surface of Caco-2 cells. J Cell Sci 114:4025–31.

29. Rawat SS, Viard M, Gallo SA, Rein A, Blumenthal R, Puri A. 2003. Modulation of entry of enveloped viruses by cholesterol and sphingolipids (Review). Mol Membr Biol 20:243–54.

30. Suomalainen M. 2002. Lipid rafts and assembly of enveloped viruses. Traffic 3:705–9.

31. Viard M, Parolini I, Sargiacomo M, Fecchi K, Ramoni C, Ablan S, Ruscetti FW, Wang JM, Blumenthal R. 2002. Role of cholesterol in human immunodeficiency virus type 1 envelope protein-mediated fusion with host cells. J Virol 76:11584–95.

32. Danthi P, Chow M. 2004. Cholesterol removal by methyl-beta-cyclodextrin inhibits poliovirus entry. J Virol 78:33–41.

33. Huang H, Li Y, Sadaoka T, Tang H, Yamamoto T, Yamanishi K, Mori Y. 2006. Human herpesvirus 6 envelope cholesterol is required for virus entry. J Gen Virol 87:277–85.

34. Medigeshi GR, Hirsch AJ, Streblow DN, Nikolich-Zugich J, Nelson JA. 2008. West Nile virus entry requires cholesterol-rich membrane microdomains and is independent of alphavbeta3 integrin. J Virol 82:5212–9.

35. Martin-Acebes MA, Gonzalez-Magaldi M, Sandvig K, Sobrino F, Armas-Portela R. 2007. Productive entry of type C foot-and-mouth disease virus into susceptible cultured cells requires clathrin and is dependent on the presence of plasma membrane cholesterol. Virology 369:105–18.

36. Diwaker D, Mishra KP, Ganju L, Singh SB. 2015. Protein disulfide isomerase mediates dengue virus entry in association with lipid rafts. Viral Immunol 28:153–60.

37. Thorp EB, Gallagher TM. 2004. Requirements for CEACAMs and cholesterol during murine coronavirus cell entry. J Virol 78:2682–92.

38. Guo H, Huang M, Yuan Q, Wei Y, Gao Y, Mao L, Gu L, Tan YW, Zhong Y, Liu D, Sun S. 2017. The Important Role of Lipid Raft-Mediated Attachment in the Infection of Cultured Cells by Coronavirus Infectious Bronchitis Virus Beaudette Strain. PLoS One 12:e0170123.

39. Inoue Y, Tanaka N, Tanaka Y, Inoue S, Morita K, Zhuang M, Hattori T, Sugamura K. 2007. Clathrin-dependent entry of severe acute respiratory syndrome coronavirus into target cells expressing ACE2 with the cytoplasmic tail deleted. J Virol 81:8722–9.

40. Wang H, Yang P, Liu K, Guo F, Zhang Y, Zhang G, Jiang C. 2008. SARS coronavirus entry into host cells through a novel clathrin- and caveolae-independent endocytic pathway. Cell Res 18:290–301.

41. Park JE, Cruz DJ, Shin HJ. 2014. Clathrin- and serine proteases-dependent uptake of porcine epidemic diarrhea virus into Vero cells. Virus Res 191:21–9.

42. Van Hamme E, Dewerchin HL, Cornelissen E, Verhasselt B, Nauwynck HJ. 2008. Clathrin- and caveolae-independent entry of feline infectious peritonitis virus in monocytes depends on dynamin. J Gen Virol 89:2147–56.

43. Pu Y, Zhang X. 2008. Mouse hepatitis virus type 2 enters cells through a clathrin-mediated endocytic pathway independent of Eps15. J Virol 82:8112–23.

44. Chu VC, McElroy LJ, Chu V, Bauman BE, Whittaker GR. 2006. The avian coronavirus infectious bronchitis virus undergoes direct low-pH-dependent fusion activation during entry into host cells. J Virol 80:3180–8.

45. McHenry EW, Reedman EJ, Sheppard M. 1938. The physiological properties of ascorbic acid: An effect upon the weights of guinea-pigs. Biochem J 32:1302–4.

46. Krzyzaniak MA, Zumstein MT, Gerez JA, Picotti P, Helenius A. 2013. Host cell entry of respiratory syncytial virus involves macropinocytosis followed by proteolytic activation of the F protein. PLoS Pathog 9:e1003309.

47. Zhu YZ, Xu QQ, Wu DG, Ren H, Zhao P, Lao WG, Wang Y, Tao QY, Qian XJ, Wei YH, Cao MM, Qi ZT. 2012. Japanese encephalitis virus enters rat neuroblastoma cells via a pH-dependent, dynamin and caveola-mediated endocytosis pathway. J Virol 86:13407–22.

48. Madu IG, Chu VC, Lee H, Regan AD, Bauman BE, Whittaker GR. 2007. Heparan sulfate is a selective attachment factor for the avian coronavirus infectious bronchitis virus beaudette. Avian Diseases 51:45–51.

49. Winter C, Schwegmann-Wessels C, Cavanagh D, Neumann U, Herrler G. 2006. Sialic acid is a receptor determinant for infection of cells by avian Infectious bronchitis virus. J Gen Virol 87:1209–16.

50. Frana MF, Behnke JN, Sturman LS, Holmes KV. 1985. Proteolytic cleavage of the E2 glycoprotein of murine coronavirus: host-dependent differences in proteolytic cleavage and cell fusion. J Virol 56:912–20.

51. Heine JW, Schnaitman CA. 1971. A method for the isolation of plasma membrane of animal cells. J Cell Biol 48:703–7.

52. Matlin KS, Reggio H, Helenius A, Simons K. 1982. Pathway of vesicular stomatitis virus entry leading to infection. J Mol Biol 156:609–31.

53. Schlegel R, Willingham M, Pastan I. 1981. Monensin blocks endocytosis of vesicular stomatitis virus. Biochem Biophys Res Commun 102:992–8.

54. Superti F, Seganti L, Ruggeri FM, Tinari A, Donelli G, Orsi N. 1987. Entry pathway of vesicular stomatitis virus into different host cells. J Gen Virol 68 ( Pt 2):387–99.

55. Marsh M, Helenius A. 2006. Virus entry: open sesame. Cell 124:729–40.

56. Martinez MG, Cordo SM, Candurra NA. 2007. Characterization of Junin arenavirus cell entry. J Gen Virol 88:1776–84.

57. Mercer J, Helenius A. 2008. Vaccinia virus uses macropinocytosis and apoptotic mimicry to enter host cells. Science 320:531–5.

58. Fretz M, Jin J, Conibere R, Penning NA, Al-Taei S, Storm G, Futaki S, Takeuchi T, Nakase I, Jones AT. 2006. Effects of Na+/H+ exchanger inhibitors on subcellular localisation of endocytic organelles and intracellular dynamics of protein transduction domains HIV-TAT peptide and octaarginine. J Control Release 116:247–54.

59. Van Leeuwen MR, Golovina EA, Dijksterhuis J. 2009. The polyene antimycotics nystatin and filipin disrupt the plasma membrane, whereas natamycin inhibits endocytosis in germinating conidia of Penicillium discolor. J Appl Microbiol 106:1908–18.

60. Vainio S, Heino S, Mansson JE, Fredman P, Kuismanen E, Vaarala O, Ikonen E. 2002. Dynamic association of human insulin receptor with lipid rafts in cells lacking caveolae. EMBO Rep 3:95–100.

61. Gallo R, Provenzano C, Carbone R, Di Fiore PP, Castellani L, Falcone G, Alema S. 1997. Regulation of the tyrosine kinase substrate Eps8 expression by growth factors, v-Src and terminal differentiation. Oncogene 15:1929–36.

62. Panda D, Rose PP, Hanna SL, Gold B, Hopkins KC, Lyde RB, Marks MS, Cherry S. 2013. Genome-wide RNAi screen identifies SEC61A and VCP as conserved regulators of Sindbis virus entry. Cell Rep 5:1737–48.

63. Henley JR, Krueger EW, Oswald BJ, McNiven MA. 1998. Dynamin-mediated internalization of caveolae. J Cell Biol 141:85–99.

64. Smith JL, Campos SK, Ozbun MA. 2007. Human papillomavirus type 31 uses a caveolin 1- and dynamin 2-mediated entry pathway for infection of human keratinocytes. J Virol 81:9922–31.

65. Cai Y, Postnikova EN, Bernbaum JG, Yu SQ, Mazur S, Deiuliis NM, Radoshitzky SR, Lackemeyer MG, McCluskey A, Robinson PJ, Haucke V, Wahl-Jensen V, Bailey AL, Lauck M, Friedrich TC, O’Connor DH, Goldberg TL, Jahrling PB, Kuhn JH. 2015. Simian hemorrhagic fever virus cell entry is dependent on CD163 and uses a clathrin-mediated endocytosis-like pathway. J Virol 89:844–56.

66. Holla P, Ahmad I, Ahmed Z, Jameel S. 2015. Hepatitis E virus enters liver cells through a dynamin-2, clathrin and membrane cholesterol-dependent pathway. Traffic 16:398–416.

67. Sun X, Yau VK, Briggs BJ, Whittaker GR. 2005. Role of clathrin-mediated endocytosis during vesicular stomatitis virus entry into host cells. Virology 338:53–60.

68. Idrissi FZ, Grotsch H, Fernandez-Golbano IM, Presciatto-Baschong C, Riezman H, Geli MI. 2008. Distinct acto/myosin-I structures associate with endocytic profiles at the plasma membrane. J Cell Biol 180:1219–32.

69. Kaksonen M, Toret CP, Drubin DG. 2005. A modular design for the clathrin- and actin-mediated endocytosis machinery. Cell 123:305–20.

70. Regan AD, Whittaker GR. 2013. Entry of rhabdoviruses into animal cells. Adv Exp Med Biol 790:167–77.

71. Piccinotti S, Kirchhausen T, Whelan SP. 2013. Uptake of rabies virus into epithelial cells by clathrin-mediated endocytosis depends upon actin. J Virol 87:11637–47.

72. Liu H, Liu Y, Liu S, Pang DW, Xiao G. 2011. Clathrin-mediated endocytosis in living host cells visualized through quantum dot labeling of infectious hematopoietic necrosis virus. J Virol 85:6252–62.

73. Mire CE, White JM, Whitt MA. 2010. A spatio-temporal analysis of matrix protein and nucleocapsid trafficking during vesicular stomatitis virus uncoating. PLoS Pathog 6:e1000994.

74. Bubb MR, Senderowicz AM, Sausville EA, Duncan KL, Korn ED. 1994. Jasplakinolide, a cytotoxic natural product, induces actin polymerization and competitively inhibits the binding of phalloidin to F-actin. J Biol Chem 269:14869–71.

75. Spector I, Braet F, Shochet NR, Bubb MR. 1999. New anti-actin drugs in the study of the organization and function of the actin cytoskeleton. Microsc Res Tech 47:18–37.

76. Ou GS, Chen ZL, Yuan M. 2002. Jasplakinolide reversibly disrupts actin filaments in suspension-cultured tobacco BY-2 cells. Protoplasma 219:168–75.

77. Sampath P, Pollard TD. 1991. Effects of cytochalasin, phalloidin, and pH on the elongation of actin filaments. Biochemistry 30:1973–80.

78. Schelhaas M, Shah B, Holzer M, Blattmann P, Kuhling L, Day PM, Schiller JT, Helenius A. 2012. Entry of human papillomavirus type 16 by actin-dependent, clathrin- and lipid raft-independent endocytosis. PLoS Pathog 8:e1002657.

79. Feng YX, Fu W, Winter AJ, Levin JG, Rein A. 1995. Multiple regions of Harvey sarcoma virus RNA can dimerize in vitro. J Virol 69:2486–90.

80. Gruenberg J. 2001. The endocytic pathway: a mosaic of domains. Nat Rev Mol Cell Biol 2:721–30.

81. Johannsdottir HK, Mancini R, Kartenbeck J, Amato L, Helenius A. 2009. Host cell factors and functions involved in vesicular stomatitis virus entry. J Virol 83:440–53.

82. Anonymous. Rab proteins as membrane organizers.

83. Stenmark H, Parton RG, Steele-Mortimer O, Lutcke A, Gruenberg J, Zerial M. 1994. Inhibition of rab5 GTPase activity stimulates membrane fusion in endocytosis. EMBO J 13:1287–96.

84. Feng Y, Press B, Wandinger-Ness A. 1995. Rab 7: an important regulator of late endocytic membrane traffic. J Cell Biol 131:1435–52.

85. Sakai T, Ohuchi M, Imai M, Mizuno T, Kawasaki K, Kuroda K, Yamashina S. 2006. Dual wavelength imaging allows analysis of membrane fusion of influenza virus inside cells. J Virol 80:2013–8.

86. Uchiyama K, Kondo H. 2005. p97/p47-Mediated biogenesis of Golgi and ER. J Biochem 137:115–9.

87. Wolf DH, Stolz A. 2012. The Cdc48 machine in endoplasmic reticulum associated protein degradation. Biochim Biophys Acta 1823:117–24.

88. Ye Y. 2006. Diverse functions with a common regulator: ubiquitin takes command of an AAA ATPase. J Struct Biol 156:29–40.

89. Cao K, Nakajima R, Meyer HH, Zheng YX. 2003. The AAA-ATPase Cdc48/p97 regulates spindle disassembly at the end of mitosis. Cell 115:355–367.

90. Ramanathan HN, Ye Y. 2012. The p97 ATPase associates with EEA1 to regulate the size of early endosomes. Cell Res 22:346–59.

91. Ritz D, Vuk M, Kirchner P, Bug M, Schutz S, Hayer A, Bremer S, Lusk C, Baloh RH, Lee H, Glatter T, Gstaiger M, Aebersold R, Weihl CC, Meyer H. 2011. Endolysosomal sorting of ubiquitylated caveolin-1 is regulated by VCP and UBXD1 and impaired by VCP disease mutations. Nat Cell Biol 13:1116–23.

92. Wong HH, Kumar P, Tay FP, Moreau D, Liu DX, Bard F. 2015. Genome-Wide Screen Reveals Valosin-Containing Protein Requirement for Coronavirus Exit from Endosomes. J Virol 89:11116–28.

93. Sieczkarski SB, Whittaker GR. 2002. Dissecting virus entry via endocytosis. J Gen Virol 83:1535–45.

94. Burkard C, Verheije MH, Wicht O, van Kasteren SI, van Kuppeveld FJ, Haagmans BL, Pelkmans L, Rottier PJ, Bosch BJ, de Haan CA. 2014. Coronavirus cell entry occurs through the endo-/lysosomal pathway in a proteolysis-dependent manner. PLoS Pathog 10:e1004502.

95. Eifart P, Ludwig K, Bottcher C, de Haan CA, Rottier PJ, Korte T, Herrmann A. 2007. Role of endocytosis and low pH in murine hepatitis virus strain A59 cell entry. J Virol 81:10758–68.

96. Nash TC, Buchmeier MJ. 1997. Entry of mouse hepatitis virus into cells by endosomal and nonendosomal pathways. Virology 233:1–8.

97. Qiu Z, Hingley ST, Simmons G, Yu C, Das Sarma J, Bates P, Weiss SR. 2006. Endosomal proteolysis by cathepsins is necessary for murine coronavirus mouse hepatitis virus type 2 spike-mediated entry. J Virol 80:5768–76.

98. Li Z, Zhao K, Lan Y, Lv X, Hu S, Guan J, Lu H, Zhang J, Shi J, Yang Y, Song D, Gao F, He W. 2017. Porcine Hemagglutinating Encephalomyelitis Virus Enters Neuro-2a Cells via Clathrin-Mediated Endocytosis in a Rab5-, Cholesterol-, and pH-Dependent Manner. J Virol 91.

99. Milewska A, Nowak P, Owczarek K, Szczepanski A, Zarebski M, Hoang-Bujnowicz A, Berniak K, Wojarski J, Zeglen S, Baster Z, Rajfur Z, Pyrc K. 2017. Entry of human coronavirus NL63 to the cell. J Virol doi:10.1128/JVI.01933-17.

100. Shirato K, Kanou K, Kawase M, Matsuyama S. 2017. Clinical Isolates of Human Coronavirus 229E Bypass the Endosome for Cell Entry. J Virol 91.

101. Pike LJ. 2009. The challenge of lipid rafts. J Lipid Res 50 Suppl:S323–8.

102. Suzuki T, Suzuki Y. 2006. Virus infection and lipid rafts. Biol Pharm Bull 29:1538–41.

103. Chazal N, Gerlier D. 2003. Virus entry, assembly, budding, and membrane rafts. Microbiol Mol Biol Rev 67:226–37, table of contents.

104. Pietiainen VM, Marjomaki V, Heino J, Hyypia T. 2005. Viral entry, lipid rafts and caveosomes. Ann Med 37:394–403.

105. Rajendran L, Simons K. 2005. Lipid rafts and membrane dynamics. J Cell Sci 118:1099–102.

106. Santoro F, Kennedy PE, Locatelli G, Malnati MS, Berger EA, Lusso P. 1999. CD46 is a cellular receptor for human herpesvirus 6. Cell 99:817–27.

107. Campbell SM, Crowe SM, Mak J. 2001. Lipid rafts and HIV-1: from viral entry to assembly of progeny virions. J Clin Virol 22:217–27.

108. Alving CR, Beck Z, Karasavva N, Matyas GR, Rao M. 2006. HIV-1, lipid rafts, and antibodies to liposomes: implications for anti-viral-neutralizing antibodies. Mol Membr Biol 23:453–65.

109. Carro AC, Damonte EB. 2013. Requirement of cholesterol in the viral envelope for dengue virus infection. Virus Res 174:78–87.

110. Vela EM, Zhang L, Colpitts TM, Davey RA, Aronson JF. 2007. Arenavirus entry occurs through a cholesterol-dependent, non-caveolar, clathrin-mediated endocytic mechanism. Virology 369:1–11.

111. Damm EM, Pelkmans L, Kartenbeck J, Mezzacasa A, Kurzchalia T, Helenius A. 2005. Clathrin- and caveolin-1-independent endocytosis: entry of simian virus 40 into cells devoid of caveolae. J Cell Biol 168:477–88.

